# Elevation-dependent patterns of snow-gum dieback are associated with subspecies’ trait differences and environmental variation

**DOI:** 10.1101/2023.12.04.569996

**Authors:** Callum Bryant, Marilyn C. Ball, Justin Borevitz, Matthew T. Brookhouse, Hannah Carle, Pia Cunningham, Mei Davey, James Davies, Ashleigh Eason, Joseph D. Erskine, Tomas I. Fuenzalida, Dmitry Grishin, Rosalie Harris, Jessica Kriticos, Aaron Midson, Adrienne B. Nicotra, Annabelle Nshuti, Jessica Ward-Jones, Yolanda Yau, Olivia Young, Helen Bothwell

## Abstract

Subalpine forests worldwide face the synergistic threats of global warming and increased biotic attack, and the collapse or transition of subalpine forests is predicted in south-eastern Australia under future climates. The recent widespread dieback of subalpine snow gum forests due to increased activity of a native wood-boring longicorn beetle suggests this process may already be underway. We investigated how variation in tree tissue traits and environmental conditions correlated with elevation-dependent spatial patterns of forest mortality.

We hypothesised that increased vulnerability of subalpine snow gums to wood-borer-mediated dieback at intermediate elevations was associated with poorly-resolved differences in traits between montane (*Eucalyptus pauciflora* subsp. *pauciflora)* and subalpine (*E. pauciflora* subsp. *niphophila)* snow-gum subspecies. We first sought to characterise variation and elevation-dependent transitions in 20 structural and drought-related functional traits among 120 healthy trees distributed along a 1000m elevation transect which spanned the subspecies transition zone. Secondly, we surveyed 774 trees across 53 sites between 1280-1980m asl. to explore associations between borer-damage severity, elevation, subspecies, and a subset of traits that differed between subspecies.

These snow-gum subspecies exhibited mean differences and/or divergent elevation responses in 25% of traits surveyed, indicating contrasting suites of traits between montane and subalpine subspecies. Increased borer-damage severity across the montane-to-subalpine subspecies transition was correlated with lower bark thickness, whereas reduced borer damage at the highest elevations was associated with greater precipitation and lower temperatures.

Our results suggest that due to possessing distinct traits associated with increased borer susceptibility, subalpine snow-gum forests will be subject to increased risk of severe borer-mediated forest dieback under warmer and drier future climates. Identifying traits contributing to species’ distribution limits and biotic-agent vulnerability remains critical for predicting, monitoring, and possibly mitigating forest and vegetation declines under future climates.

## INTRODUCTION

Global climate change is challenging many forested ecosystems with elevated temperatures and increasingly severe and frequent drought, heat waves, and outbreaks of biotic agents (Meehl and Tebaldi, 2004; Chou *et al*., 2013; Davy *et al*., 2017; Pureswaran, Roques and Battisti, 2018; Harvey *et al*., 2020; IPCC, 2022). Escalating forest dieback occurrences in response to changing abiotic and biotic pressures have significant implications for ecosystem structure, biodiversity, and terrestrial fluxes of carbon and water (Allen, Breshears and McDowell, 2015; Trumbore, Brando and Hartmann, 2015; Clark *et al*., 2016). The leading hypothesised mechanisms of forest decline under global change revolve around carbon and hydraulic deficits in tree tissues and their interactions with the intensity, duration and frequency of disturbance (McDowell *et al*., 2022). Characterising the resilience of many unmanaged forests to these challenges remains challenging due to a limited understanding of the key biotic elements present, factors influencing population dynamics in multi-trophic interactions, and divergent responses of co-occurring species to changing conditions (Fei *et al*., 2017; Seidl *et al*., 2017; Anderegg *et al*., 2018). Thus, characterising the differential vulnerabilities of native forest species to agents of mortality is critical to improving our ability to predict, monitor, and possibly mitigate declines of native forests under future climates (Millar and Stephenson, 2015).

While forest dieback and vegetation decline have been observed across all forested biomes (Allen *et al*., 2010), montane and subalpine forests are among the most sensitive to increased temperatures (Hammond *et al*., 2022; IPCC, 2022). Increases in mean temperatures may be further enhanced by elevation-dependent warming occurring near the 0°C isotherm caused by snow-ice albedo feedbacks with temperature and radiation fluxes, a phenomenon observed across the majority of mountain regions (Pepin and Lundquist, 2008; Pepin *et al*., 2015). As warmer temperatures often support increased insect growth, reproduction, locomotion, dispersal and survival (Huey *et al*., 2012; Kingsolver, Higgins and Augustine, 2015; Roitberg and Mangel, 2016; González-Tokman *et al*., 2020), the interactive effects of changing temperatures on insect population dynamics and increased access to previously cold protected forests threaten montane-subalpine forest systems with increased pressure from biotic agents (Veblen *et al*., 1991; Gan, 2004; Hicke *et al*., 2006; Mantgem *et al*., 2009; Cudmore *et al*., 2010; Reed and Hood, 2021).

In south-eastern Australia, the collapse or transition of high-elevation forests under future climates has been predicted with high probability due to the effects of warming on snow cover and fire frequency on subalpine specialist or fire-sensitive species (IPCC, 2022; Lawrence *et al*., 2022; Verrall *et al*., 2023). However, recent observations of dieback of subalpine *Eucalyptus pauciflora* Sieber ex Spreng woodlands suggests that increased activity of a native, wood-boring long-horn beetle, *Phoracantha mastersii* (Pascoe) is also an emergent proximate threat (Elliott, Ohmart and Wylie, 1998; Ward-Jones, 2020; Meyer, 2021; Brookhouse, Ward-Jones and Meyer, 2022). Severe forest damage due to *Phoracantha* beetles (Coleoptera: Cerambycidae) has been reported in several *Eucalyptus* forests (Hanks, Paine and Millar, 1991, 2005; Caldeira *et al*., 2002; Paine, Steinbauer and Lawson, 2011; Nahrung *et al*., 2014; Seaton *et al*., 2015; Seaton, Matusick and Hardy, 2020). *Phoracantha* larvae feed circumferentially on the cambial zone of living trees, causing increasing hydraulic and metabolic dysfunction, resulting in canopy decline and ultimately stem death (Wang, 1995; Brookhouse, Ward-Jones and Meyer, 2022).

In south-eastern Australia, an emergent borer-mediated woodland snow-gum dieback was reported by National Park rangers as early as 2017, and observations of dieback have since been recorded in several high-elevation National Parks along the great-dividing range spanning south-eastern Australia (Brookhouse, Ward-Jones and Meyer, 2022). While the spatial and temporal dynamics of the current snow-gum dieback remain poorly resolved, preliminary observations suggest increased borer activity has differentially impacted several snow-gum species and subspecies: *Eucalyptus pauciflora* subsp. *niphophila*, *E. pauciflora* subsp. *debeuzevillei* and *Eucalyptus lacrimans* (Ward-Jones, 2020; Brookhouse, Ward-Jones and Meyer, 2022). Surveys in *E. pauciflora subsp. niphophila* at elevations between 1600-1950m found borer damage to be negatively correlated with elevation, positively correlated with steeper slopes and larger diameter stems, however not influenced by stand-density or aspect (Ward-Jones, 2020). Further, while not captured in these surveys, snow-gum dieback appears to decline sharply below 1600m corresponding approximately with the transition between subalpine and montane snow-gum forests (Slee *et al*., 2015).

In other forest systems, *Phoracantha* outbreaks have been associated with extreme drought conditions, and larval penetration of host bark to be negatively correlated with plant tissue water content (Hanks, Paine and Millar, 1991; Nahrung *et al*., 2014; Seaton *et al*., 2015; Seaton, Matusick and Hardy, 2020). As drought conditions often involve the co-occurrence of tree water deficits and elevated temperatures, the relative influences of increased tree vulnerability due to drought-related water stress and amplified insect activity due to temperature increases are not easily distinguished (Seidl *et al*., 2017). As precipitation and temperature are inversely correlated along elevation transects in south-eastern Australia (Körner and Cochrane, 1985; Körner, 2007), the ultimate causes of current elevation-dependent spatial patterns of snow-gum dieback may be influenced by moisture and temperature variation, the limits of snow-gum subspecies’ distributions and a warming climate.

In mountain systems, intraspecific trait variation with increasing elevation is expected in response to declining temperatures (Fajardo and Piper, 2011; Mayor *et al*., 2017; Körner, 2021); however, poorly resolved transitions between discrete populations or subspecies can hinder these predictions. In south-eastern Australia, the vegetation transition from montane to subalpine woodlands generally coincides with a transition from a lowland-montane distributed snow-gum subspecies, *E. pauciflora* subsp. *pauciflora* (Johnson and Blaxell, 1973; Slee *et al*., 2015), to stands of one of several geographically isolated subalpine snow-gum subspecies (Slee *et al*., 2015). Currently, within the *E. pauciflora* subsp. complex, four geographically distinct high-elevation snow-gum subspecies are described (*E. pauciflora* subsp. *acerina, debeuzevillei, hedraia and niphophila*). However, structural and functional differences among snow-gum subspecies that may impact differential wood borer vulnerability and drought resilience remain poorly resolved.

Several studies have investigated morphological and physiological variation along elevation transects without differentiating between snow-gum subspecies, with transects spanning montane-distributed *Eucalyptus pauciflora* subsp*. pauciflora* to one of two subalpine-distributed snow gums, either *E. pauciflora* subsp. *debeuzevillei* (Pryor, 1956; Green, 1969) or *E. pauciflora* subsp. *niphophila* (Slatyer, 1977c, 1977a, 1977c; Slatyer and Ferrar, 1977a, 1977b; Slatyer, 1978; Körner and Cochrane, 1985; O’sullivan *et al*., 2013). In field and progeny greenhouse experiments, these studies have captured elevation-associated variation in leaf morphology, bark thickness, germination and growth rates and vulnerability to frost-kill (Pryor, 1956; Green, 1969), photosynthetic temperature optima and its seasonal acclimation (Slatyer, 1977a, 1977c, 1977b, 1978; Slatyer and Ferrar, 1977b, 1977a), seasonal acclimation of leaf respiration and photosynthetic heat tolerance (O’sullivan *et al*., 2013) and summer variation in diurnal patterns of stomatal conductance and leaf water relations (Körner and Cochrane, 1985). However, due to the specific traits studied and limited resolution of variation within subspecies’ respective elevation distributions, these studies afford limited resolution of mean trait differences between snow-gum subspecies that may contribute to the observed elevation-dependent patterns of borer dieback.

Resolving the phylogenetic underpinning of spatial patterns of train variation is important because tree resilience to *Phoracantha* outbreaks can be with variation in structural traits such as stem size and tissue allocations (bark thickness and tissue densities) and their interaction with tissue water content (Nahrung *et al*., 2014; Seaton, Matusick and Hardy, 2020). Snow gums have been observed to maintain relatively stable tissue water potentials by closing stomata early in the day to avoid water loss, as well as show minimal osmotic adjustment during summer (Körner and Cochrane, 1985). However, little is known about variation in other traits required to maintain a positive water balance during drought, such as sapwood-to-leaf area ratios, tissue water storage and hydraulic capacitance, background cuticular water losses and maintenance of hydraulic conductance (Mencuccini *et al*., 2019; McDowell *et al*., 2022). As the lower elevation limits of borer-mediated dieback appear to be associated with the elevation-associated transition between snow-gum subspecies (Ward-Jones, 2020), resolving subspecies differences in these structural and drought-related functional traits is critical to understanding differential vulnerability, modelling future borer spread and management for biodiversity.

The present study aimed to identify patterns of trait and environmental variation in the montane-subalpine landscape that may contribute to increased borer damage observed at intermediate elevations. Observations of sharp increases in borer damage at intermediate elevations led us to question previous conclusions in the literature that snow-gum traits vary continuously with elevation across the transition from *Eucalyptus pauciflora* subsp. *pauciflora* and *E. pauciflora* subsp. *niphophila.* We first conducted an exploratory survey of variation in structural and drought-related functional traits among snow-gum subspecies and in response to elevation in 120 healthy trees distributed along a single 1000m elevation transect spanning the subspecies’ transition zone. Twenty traits were characterised: leaf size; Huber value (sapwood to leaf area ratio); leaf mass per unit area; bole and branch sapwood density; leaf and stem saturated water content and hydraulic capacitance; leaf and stem minimal surface conductance; bole and branch relative bark thickness; bole bark density; xylem vessel diameter and density, xylem lumen and fibre fractional areas, fibre density and theoretical hydraulic conductance of vessels and stems. We hypothesised that trait values would differ stepwise across the subspecies transition zone and that trait variation in response to environmental change along an elevation transect may differ between snow-gum subspecies. We then conducted a second survey, including healthy and dieback-affected individuals, to assess associations between increased borer-damage severity and a subset of traits based on the first analysis, particularly wood density and bark thickness. Here we hypothesised that differences in wood density and bark thickness may be associated with the severity of borer damage in each snow-gum subspecies. This second survey also explored associations between spatial variation in environmental variables and borer-damage severity.

## METHODS

### Study site, subspecies and longitudinal climate trends

This study was conducted within Kosciuszko National Park (KNP), New South Wales (NSW) (148.358, -36.431; Figure **1a**). In KNP, *Eucalyptus. pauciflora* subsp. *pauciflora* occurs in pure and mixed stands up to ∼1550m; above ∼1650m *E. pauciflora* subsp. *niphophila* occurs in pure stands, with subspecies distinguishable by leaf and bud characteristics (Slee *et al*., 2015). Elevations in the transition zone, ∼1550-1650m (depending on aspect and microclimate), contain trees with mixed *subsp. pauciflora* - *subsp. niphophila* leaf and bud characteristics (Figure **1b**). Preliminary field surveys and visual observations showed strong elevation-dependent variation in forest decline and evidence of borer damage (Figure **1c,d**)(Ward-Jones, 2020). Kosciuszko National Park is located within the Australian Alpine Bioregion and is dominated by a montane climate. As changing regional climate conditions are predicted to challenge subalpine communities, we describe regional warming via longitudinal records of maximum snow depth and snow cover from 1955-2020 for three sites differing in elevation within KNP (1430m, 1620m, 1830m) (https://www.snowyhydro.com.au/generation/live-data/snow-depths/). These records demonstrate persistent longitudinal reductions in the maximum depth of snow cover consistent across all elevations (-0.541 ± 0.174 cm year^-1-^; Figure **1e**), and reductions in the duration of annual snow cover due to a progressively earlier end of the snow season (-0.373 ± 0.060 days year^-1^; Figure **1f**). Longitudinal records of regional drought conditions for the study region were illustrated using the standardised precipitation-evapotranspiration index (SPEI) values obtained from the SPEI global drought monitor (- 36.25, 148.25, 1955-2020; Figure **1g**, https://spei.csic.es; however, see also Appendix S1: *Supplementary methods* and Figure **S1**).

**FIGURE 1.**
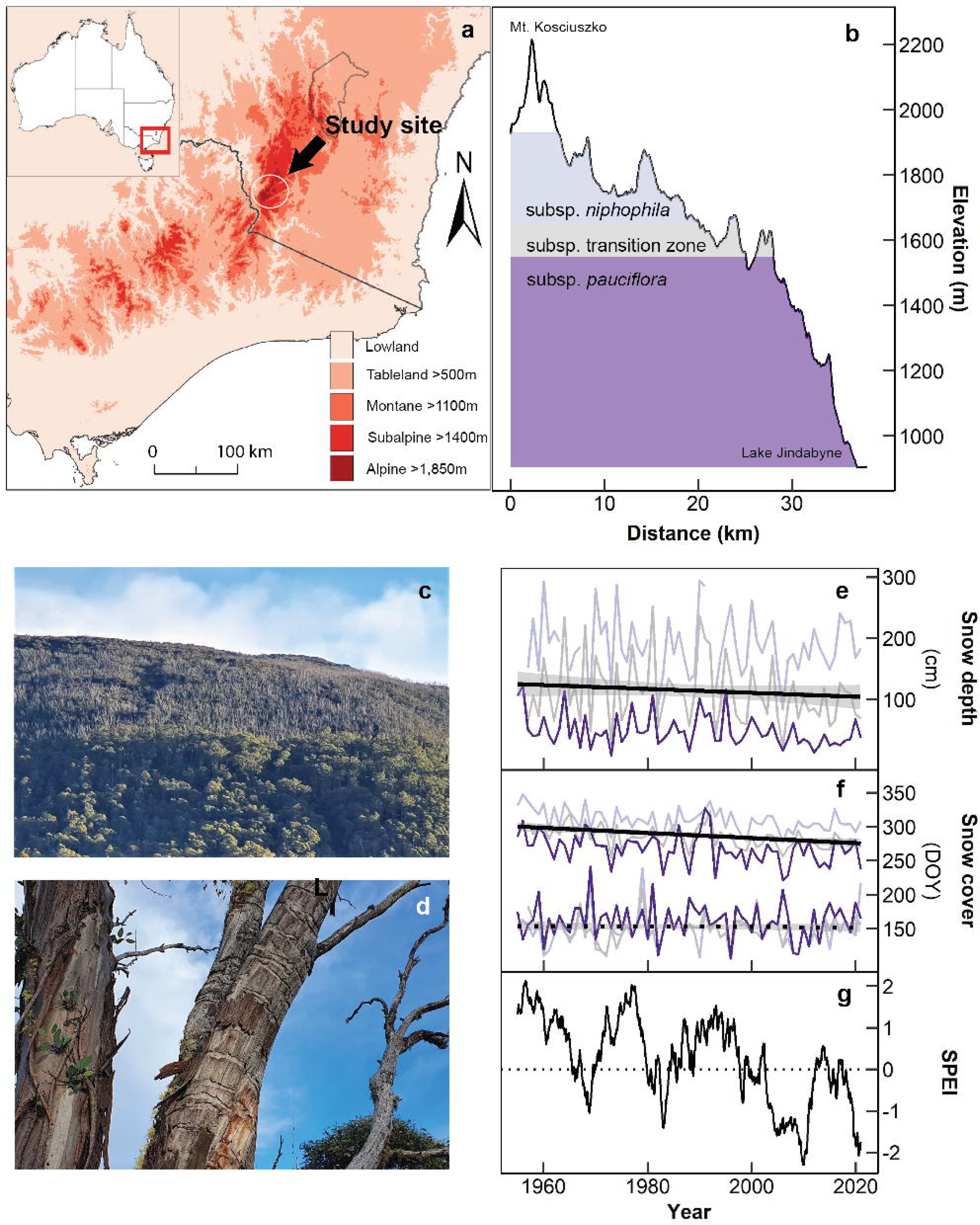
**(a)** Location of subalpine study region within 90m relief map of south-eastern Australia. **(b)** Elevation profile (Perisher Valley) illustrating approximate elevations associated with the transition from *Eucalyptus pauciflora subsp. pauciflora* (∼<1550m) *to E. pauciflora subsp. niphophila* (∼>1650m). **(c-d)** Images of the study region illustrating extensive forest dieback and associated evidence of wood borer galleries. **(e)** Time series data (1955-2020) from three elevations, 1830m (pale), 1620m (grey), 1430m (dark), depicting longitudinal declines in maximum snow depths (Linear mixed model ANOVA: (*F*_1,195_=9.63, *P*<0.01) **(f)** Longitudinal shortening of annual snow cover period due to earlier end to the snow season (solid line, Linear mixed model ANOVA (*F*_1,195_=38.2, *P*<.00001) rather than changes in season onset (dotted line; not significant). **(g)** Inter-decadal trend (1955-2020) toward longer and more intense periods of regional drying, illustrated via increasingly negative standardised precipitation-evapotranspiration index (SPEI, 24-month rolling average), a drought index calculated as the difference between monthly precipitation and potential evapotranspiration (Vicente-Serrano et al. 2010).

### Trait variation between subspecies and in response to elevation

Variation in twenty functional traits between subspecies and in response to elevation was surveyed in 120 trees distributed along a single east-west elevation transect in the Perisher Valley, NSW (Appendix S1: Figure S**2**). Traits of ten healthy trees were surveyed at each of 12 sites spanning a total distance of ∼31 km between 920-1930m. *E. pauciflora* subsp. *pauciflora* was surveyed across six sites (*n*=60), *subsp. niphophila* was surveyed across five sites (*n*=50), with one site included in the subsp. transition zone (*n*=10). At each site, ten living trees were selected by walking on contour and sampling at ∼25m intervals. Trees were geotagged and photographed to enable the cataloguing of traits for specific individuals over iterative field campaigns conducted within a five-week period between late December 2021 and early January 2022. Environmental data for each site are summarised in *Supplementary Datasheet 2*.

Traits surveyed captured variation across several aboveground organs; leaves, canopy branch stem tissues, and the tree bole. Leaf and branch traits were characterised in tissues obtained from sun-exposed canopy branches up to 1m long. Bole traits were determined at breast height in the largest stem. Variation in structure and tissue organisation was explored via leaf size, sapwood-to-leaf area ratios (Huber value; cm^2^ m^-2^), leaf mass per unit leaf area (LMA; g m^-2^) and relative bark thickness in the bole (RBT_bole;_ % of diameter) and the branch (RBT_branch_). Variation in tissue density (g cm^-3^) was explored in bole bark (BD_bark_), bole sapwood (WD_bole_), and branch sapwood (WD_branch_). Variation in tissue water storage, ease of water release and background water loss was measured via saturated water content in leaves (SWC_leaf_; g_water_ g^-1^_dry mass_ at 0MPa) and stems (SWC_stem_), hydraulic capacitance in leaves (*C*_leaf_; g_water_ g^-1^_dry mass_ MPa^-1^) and stems (*C*_stem_), and minimum surface conductance to water vapour in leaves (*g*_min leaf_; mmol_water_ m^-2^ s^-1^) and stems (*g*_min stem_). Additionally, we explored variation in several hydraulically related traits derived from microscopic analyses of branch sapwood cross sections. These included xylem vessel diameter (µm), xylem vessel density (vessel no. mm^-2^), xylem lumen fraction of total sapwood area (lumen fraction; %), branch sapwood fibre density, i.e. density of total sapwood area - xylem lumen fraction (fibre density; g cm^-3^) and theoretical hydraulic conductivity of a mean vessel (*K*v; mg_water_ m^-1^ s^-1^ MPa^-1^) and per stem cross-sectional area (*K*s; kg_water_ m^-1^ s^-1^ MPa^-1^). Detailed protocols used to determine each trait are provided in the *Supplementary methods*.

### Associations between traits, environmental variation and borer-mediated damage

To assess asociations between borer-induced tree dieback, environment and a subset of traits, we surveyed 774 trees from 53 sites for borer-damage severity, diameter at breast height (DBH), bark thickness (BT) and wood density (WD) in May 2022. Sites were selected at ∼80m elevation intervals spanning an elevation gradient from 1270-1970m, which captures the montane-subalpine-alpine vegetation zones and the transition between the distributions of snow-gum subspecies (Appendix S1: Figure S**2**). Where possible, sites were selected at each elevation to capture a spectrum of aspects and stand conditions. As this study was primarily focused on the associations between borer-induced dieback, elevation and subspecies traits, we attempted to avoid the confounding effects of recent fire damage by only selecting sites with no immediate history of fire-killed stems, i.e. no dead stems lacking evidence of borer damage. Sites included in the study spanned 38.0 km E-W and 31.1 km N-S (Appendix S1: Figure S**2**).

At each site, 15 living trees were selected by walking on contour and sampling at ∼25m intervals. As snow gums can take a multi-stemmed (mallee) growth form depending on fire history, for each individual, borer damage on the largest stem was scored using a previously established (0-4) borer-damage severity index (Ward-Jones, 2020): where 0 = healthy; 1= circumferentially continuous live bark with visible borer frass holes and wounding stains, 2 = patchy live bark/ patchy dead bark, 3 = dead stem (bark still attached), 4 = dead stem (bark fallen away) (Figure **2**). On the largest stem, DBH, bark thickness and wood density were also measured. Bark thickness (BT) was averaged from three measurements made at 120° at breast height using a bark depth gauge (Model 4340. Keuffel & Esser Co., Hoboken, N.J.). Relative bark thickness was calculated as two-sided bark thickness divided by DBH (RBT = ((2 X BT)/DBH)*100)(Rosell *et al*., 2014). Bole sapwood wood density (g cm^-3^) was assessed using ∼1 cm^3^ samples taken at breast height, perpendicular to the stem’s lean/ aspect, either with a slide hammer corer or chisel (Pérez-Harguindeguy *et al*., 2013). Samples were stored in falcon tubes humidified with moistened tissues to minimise changes in volume due to dehydration between sampling and measurement. Sample volume was measured by the displaced water volume using a precision balance (XP 205 Metter Toledo balance, Mettler – Toledo – Ltd., Griefense, Switzerland), and dry mass was determined after > 72hr oven drying at 70 °C. GPS coordinates were recorded for each tree at the time of sampling (ArcGIS Survey 123, ESRI) to enable the exploration of associations between borer-damage severity and environmental parameters associated with elevation. Modelled soil variables (soil depth to bedrock (m); clay (%), silt (%) and sand (%); pH and soil organic content (tons/hectare)) were obtained from the *SoilGrids* 250m global dataset (Hengl *et al*., 2017). Site bioclimatic variables at 1km resolution were obtained from *Worldclim 2.1* (Fick and Hijmans, 2017).

**FIGURE 2.**
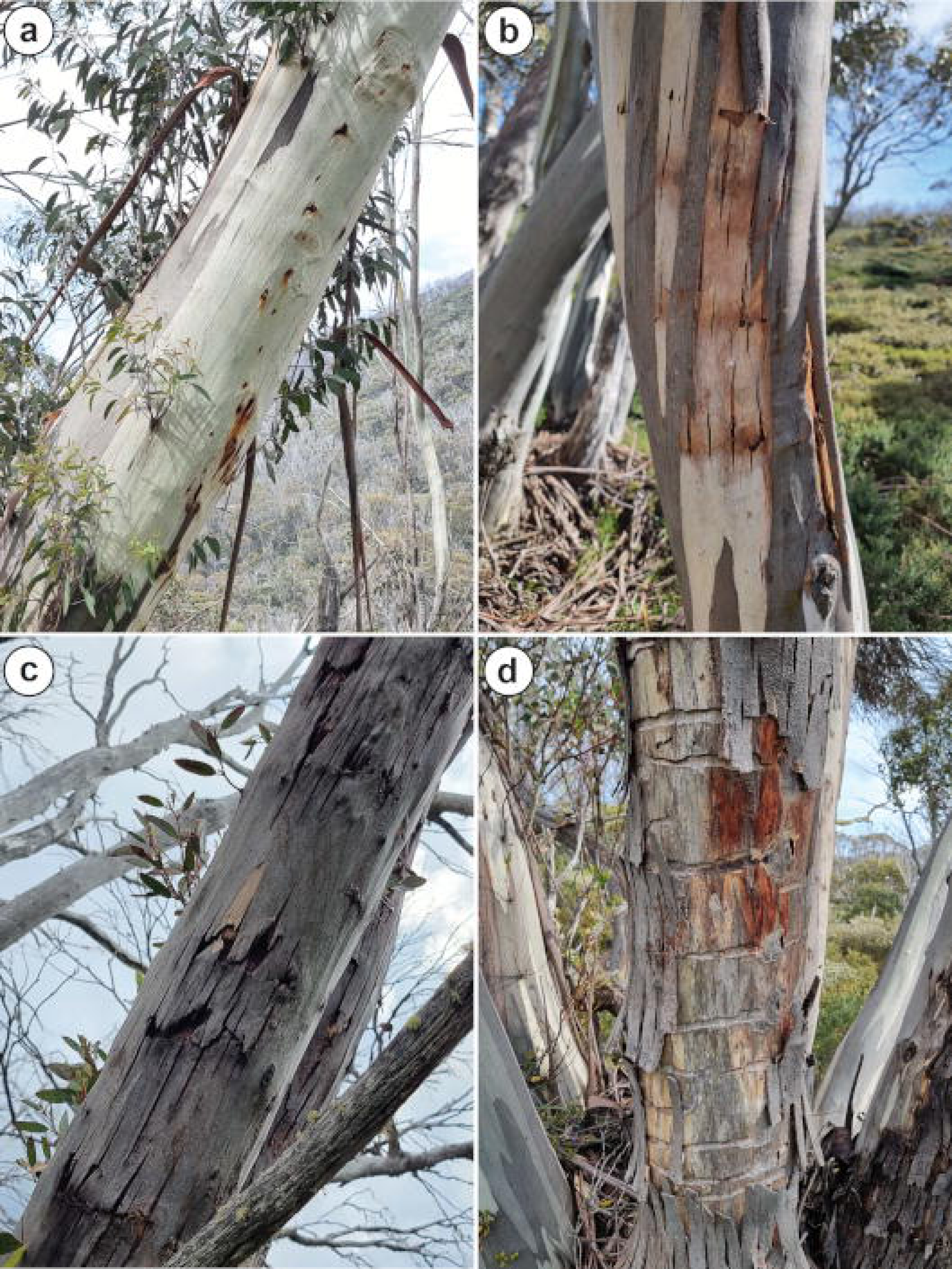
Wood-borer damage severity was scored using a (0-4) damage severity index (Ward-Jones, 2020), where: 0 = healthy; 1= circumferentially continuous live bark with visible borer frass holes and wounding stains **(a)**; 2 = patchy live bark/ patchy dead bark **(b)**; 3 = dead stem (bark still attached) **(c)**; 4 = dead stem (bark fallen away) **(d)**. *Note:* to be included in the stud, individuals with scores 3 and 4 also had to have live bark at breast height.

### Analyses

All analyses were conducted using R version 4.2.1 (R Core Team, 2022). Regional longitudinal records of snow depth and snow cover duration (beginning and end) were analysed with linear mixed models with elevation as a random factor. Mean subspecies traits across elevations were summarised using the _EMMEANS_ package, with site included as a random effect (Searle, Speed and Milliken, 1980). To discern whether traits that differed between subspecies varied continuously with elevation or shifted sharply between subspecies, trait variation was analysed with linear models with each trait as the response variable and elevation, subspecies, aspect and DBH as fixed effects. If both elevation and subspecies were significant, the models were rerun with an interaction term; if aspect and DBH were significant, they were included as random factors using the _LMER_ package (Bates *et al*., 2015). A Holm-Bonferroni correction was applied across trait comparison *P*-value outputs to minimise the potential for Type 1 errors arising from multiple comparisons (Holm, 1979). Multi-dimensional trait clustering and coordination within subspecies were explored with principal component analyses (PCA) after Z-score normalisation to enable equal weighting of predictors using the _FACTOMINER_ package (Lê, Josse and Husson, 2008). Values given throughout the text are mean ± SE unless otherwise stated.

As we scored dieback damage using an ordinal borer-damage severity index, proportional odds models (cumulative link models) were used to determine whether a given predictor variable (either a grouping or a continuous variable) influenced the probabilities of observations falling into a higher borer-damage severity category. Differences between subspecies in mean trait values were again characterised using the larger, more robust data set from the second survey (i.e. *n* = 774 rather than *n* = 120). Allometric relationships between RBT and DBH were also characterised to compare bark thicknesses between subspecies across all stem sizes. Cumulative link models were constructed using the clmm() and clm() functions in the _ORDINAL_ package in R (Agresti, 2010; Christensen, 2015, 2016); a logit link function was used after comparing with other link functions using likelihood ratio tests. Continuous predictors were rescaled using Z-score transformation to enable comparison of their relative contributions to borer-damage predictions. Initially, we used a mixed model with subspecies, elevation, WD, DBH, BT, RBT, and aspect treated as fixed effects and site as a random effect. However, to enable the inclusion of a subspecies × elevation interaction term, we then repeated this model with only fixed effects. The goodness of model fit for ordinal logistic regression (*R*^2^_pseudo_) was estimated using the Nagelkerke (Craig and Uhler) method in the _RCOMPANION_ package (Nagelkerke, 1991). Given minimal borer damage in subsp. *pauciflora,* we further explored the relationship between declining borer-damage severity and increasing elevation by repeating the above mixed ordinal model using subsp. *niphophila* data only. Due to the multi-collinearity of elevation and many environmental variables (Appendix S1: Figure S**3**) and low spatial resolution of modelled environmental data (Table S3), a comparison of the relative predictive strength of environmental variables within a single ordinal regression model was not possible. Therefore, to further probe the decline in borer-damage severity with increasing elevation observed in subsp. *niphophila,* we explored the predictive strength of modelled environmental variables highly correlated with elevation (*r*^2^>.7, Appendix S1: Figure S**3**) via comparison of several models containing single-term elevation substitutions, using likelihood ratio statistics.

## RESULTS

### Trait variation between subspecies and in response to elevation

Of the 20 structural and drought-related functional traits surveyed across leaf, stem and bole tissues of healthy trees, ten traits (50%) differed between subspecies when averaged across all elevations (Table **1**). Underlying these bulk subspecies differences, we observed continuous variation associated with elevation, differences in mean trait values, as well as differences in trait variation in response to elevation between subspecies (Figure **3**, Appendix S1: Table S**1**,**2**). Four traits (20% of surveyed traits) varied continuously across the elevation transect; Huber values (Figure **3b**) and LMA (Figure **3c**) increased with elevation, whereas vessel-specific hydraulic conductivity (*K*v, Figure **3i**) and stem-specific hydraulic conductivity (*K*s, Figure **3j**) decreased with elevation. Four traits (20% of surveyed traits) differed between subspecies but varied little within each subspecies elevation range. Relative to subsp. *pauciflora*, subsp. *niphophila* had smaller mature leaves (F_1,97_=104.58; *P*<.0001; Figure **3a**), lower bole relative bark thickness RBT_bole_ (*F*_1,50_=31.4, *P*<.05; Figure **3d**), higher branch relative bark thickness, and smaller xylem vessel diameters (*F*_1,57_=14.1, *P*<.001, Figure **3f**). We observed a subspecies × elevation interaction in a single trait, xylem lumen fraction, which declined with elevation in subsp. *pauciflora* but increased with elevation in subsp. *niphophila* (*F*_1,102_ = 9.65, *P* <.01, Figure **3g**).

**FIGURE 3.**
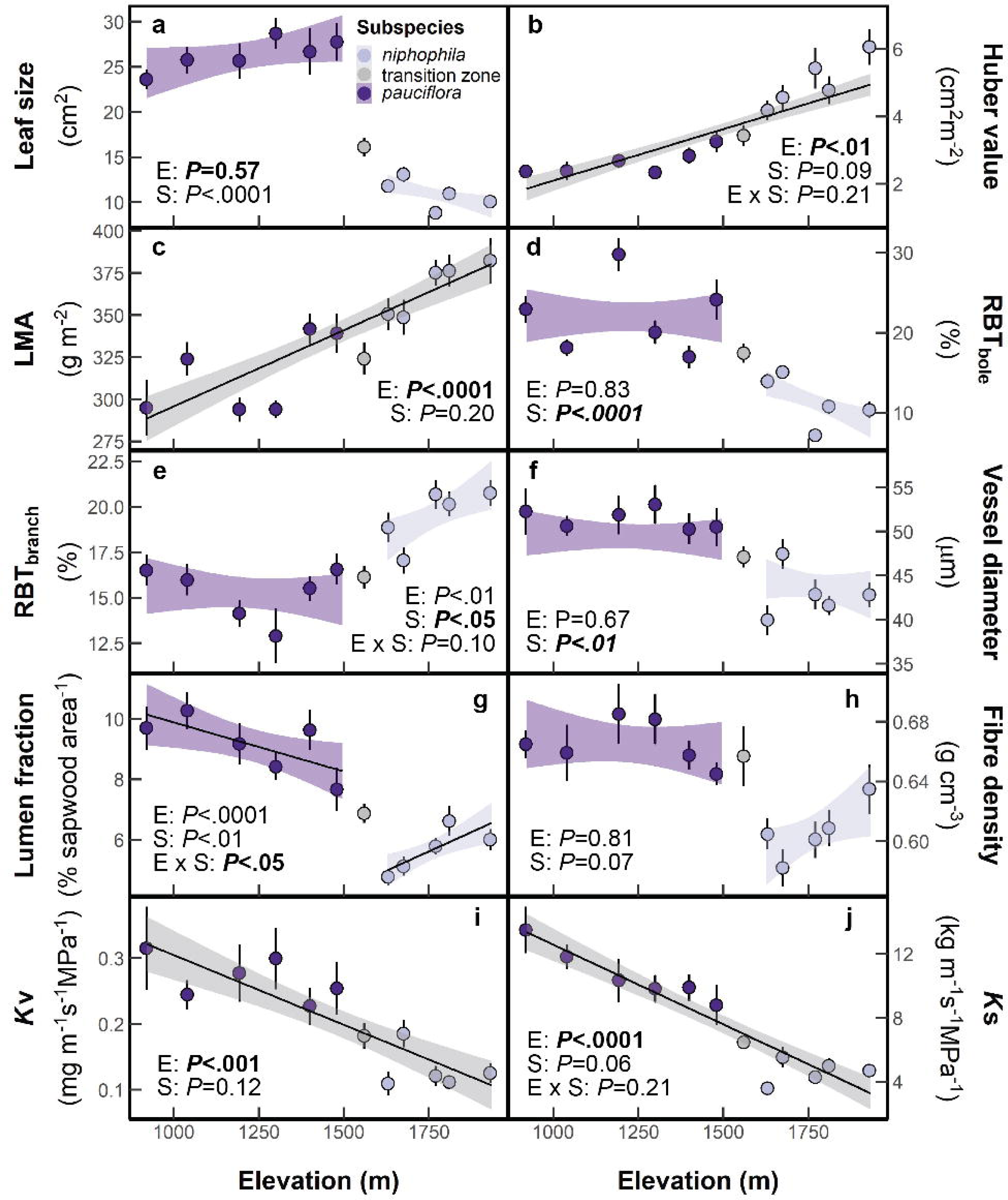
Variation in structural and tissue traits along an elevation transect spanning distributions of *Eucalyptus pauciflora* subsp. *pauciflora* (dark) and *E. pauciflora* subsp. *niphophila* (pale): **(a)** Mature leaf size; **(b)** sapwood to leaf area ratio (Huber value; HV); **(c)** leaf mass per unit area (LMA); **(d, e)** bole and branch relative bark thickness (RBT, % of diameter); **(f)** Xylem vessel diameter, **(g, h)** xylem lumen fraction and density of fiber fractions; **(i)** vessel specific hydraulic conductivity (*K*v) and **(j)** stem specific hydraulic conductivity (*K*s). Points are site means ± SE. Solid lines denote significant (*P <* .05) trait variation across elevation, as derived from linear mixed models (SI Table 1, 2, *n* = 120). Shaded regions depict 95% confidence intervals on the regression. *P*-values were corrected for multiple comparisons using the Holm-Bonferroni method.

**TABLE 1.** Comparison of mean structural and drought-related functional leaf, branch and bole traits surveyed in *Eucalyptus pauciflora* subsp. *pauciflora* and subsp. *niphophila* (n = 57-60 and 48-50, respectively). Values are marginal means ± SE averaged across all elevations with site as a random effect. *P*-values are corrected for multiple comparisons using the Holm-Bonferroni method.

| Trait | Abbrev. | Units | *E. pauciflora* subsp. |  | *P*-value |
| --- | --- | --- | --- | --- | --- |
|  |  |  | *pauciflora* | *niphophila* |  |
| Tree height | - | m | 9.68 ± 3.30sd | 6.36 ± 2.75sd | 0.14 |
| Leaf size | LS | cm<sup>2</sup> | 26.3 ± 0.7 | 10.9 ± 0.8 | <.0001 |
| Huber value (sapwood: leaf area ratio) | HV | cm<sup>2</sup> m<sup>-2</sup> | 2.65 ± 0.23 | 5.00 ± 0.25 | <.01 |
| Leaf mass per unit area | LMA | g m<sup>-2</sup> | 315 ± 8 | 367 ± 9 | <.05 |
| Wood density | WD<sub>branch</sub> | g cm<sup>-3</sup> | 0.606 ± 0.007 | 0.571 ± 0.007 | .052 |
| | WD<sub>bole</sub> | g cm<sup>-3</sup> | 0.748 ± 0.012 | 0.706 ± 0.015 | 0.34 |
| Bark thickness | RBT<sub>branch</sub> | % of diam. | 15.3 ± 0.6 | 19.5 ± 0.7 | <.01 |
| | RBT<sub>bole</sub> | % of DBH. | 22.0 ± 1.7 | 11.4 ± 1.8 | <.05 |
| Bark density | BD<sub>bole</sub> | g cm<sup>-3</sup> | 0.452 ± 0.015 | 0.375 ± 0.016 | 0.057 |
| Saturated water content | SWC<sub>leaf</sub> | g<sub>water</sub> g<sup>-1</sup> dry mass | 1.23 ± 0.03 | 1.10 ± 0.03 | 0.052 |
| | SWC<sub>branch</sub> | g<sub>water</sub> g<sup>-1</sup> dry mass | 1.05 ± 0.02 | 1.08 ± 0.02 | 0.11 |
| Minimum surface water vapour conductance | g<sub>min</sub> leaf | mmol m<sup>-2</sup> s<sup>-1</sup> | 10.23 ± 0.90 | 6.46 ± 0.99 | 0.14 |
| | g<sub>min</sub> stem | mmol m<sup>-2</sup> s<sup>-1</sup> | 15.9 ± 1.8 | 13.3 ± 2.0 | 0.11 |
| Hydraulic capacitance | C<sub>leaf</sub> | g<sub>water</sub> g<sup>-1</sup> dry mass MPa<sup>-1</sup> | 0.0569 ± 0.0021 | 0.0456 ± 0.0023 | .052 |
| | C<sub>branch</sub> | g<sub>water</sub> g<sup>-1</sup> dry mass MPa<sup>-1</sup> | 0.105 ± 0.006 | 0.0881 ± 0.007 | 0.11 |
| Vessel diameter | VD | μm | 51.4 ± 0.9 | 42.9 ± 0.9 | <.01 |
| Vessel density | VMM | no. vessel mm<sup>-2</sup> | 41.4 ± 2.6 | 36.9 ± 2.9 | .11 |
| Lumen fraction | LF | % per cross-sectional area | 9.14 ± 0.35 | 5.65 ± 0.39 | <.01 |
| Fibre fraction density | FD | g cm<sup>-3</sup> | 0.665 ± 0.007 | 0.605 ± 0.007 | <.01 |
| Vessel-specific hydraulic conductivity | Kv | mg m<sup>-1</sup> s<sup>-1</sup> MPa<sup>-1</sup> | 0.268 ± 0.0134 | 0.131 ± 0.015 | <.0001 |
| Stem-specific hydraulic conductivity | Ks | kg m<sup>-1</sup> s<sup>-1</sup> MPa<sup>-1</sup> | 10.68 ± 0.55 | 4.59 ± 0.60 | <.001 |

**TABLE 2.**
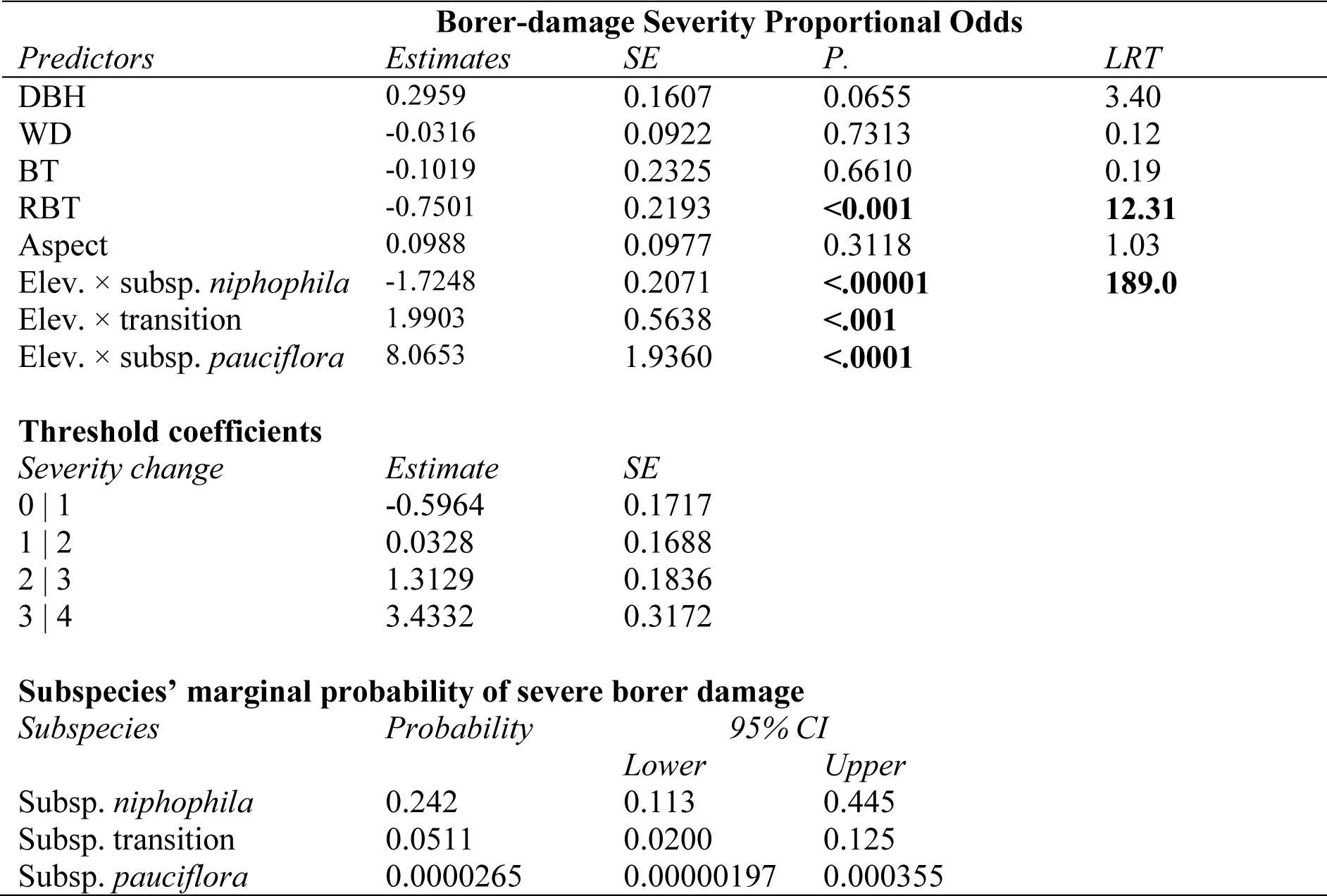
Borer-damage severity (0-4) proportional-odds cumulative logit model parameter estimates. Note coefficient estimates and SE are log-odds of a shift in the borer-damage severity category (0-4) per unit SD shift in the predictor variable. Predictors are scaled and centred; therefore, estimates are best interpreted as the relative predictor strength and direction of the association. *P* values were obtained from likelihood-ratio tests (LRT) of explanatory variables while controlling for remaining predictors. *n* = 774.

| **Borer-damage Severity Proportional Odds** |  |  |  |  |
| --- | --- | --- | --- | --- |
| *Predictors* | *Estimates* | *SE* | *P.* | *LRT* |
| DBH | 0.2959 | 0.1607 | 0.0655 | 3.40 |
| WD | -0.0316 | 0.0922 | 0.7313 | 0.12 |
| BT | -0.1019 | 0.2325 | 0.6610 | 0.19 |
| RBT | -0.7501 | 0.2193 | **<0.001** | **12.31** |
| Aspect | 0.0988 | 0.0977 | 0.3118 | 1.03 |
| Elev. × subsp. *niphophila* | -1.7248 | 0.2071 | **<.00001** | **189.0** |
| Elev. × transition | 1.9903 | 0.5638 | **<.001** |  |
| Elev. × subsp. *pauciflora* | 8.0653 | 1.9360 | **<.0001** |  |
| **Threshold coefficients** |  |  |  |  |
| *Severity change* | *Estimate* | *SE* |  |  |
| 0 \| 1 | -0.5964 | 0.1717 |  |  |
| 1 \| 2 | 0.0328 | 0.1688 |  |  |
| 2 \| 3 | 1.3129 | 0.1836 |  |  |
| 3 \| 4 | 3.4332 | 0.3172 |  |  |
| **Subspecies' marginal probability of severe borer damage** |  |  |  |  |
| *Subspecies* | *Probability* | *95% CI* |  |  |
|  |  | *Lower* | *Upper* |  |
| Subsp. *niphophila* | 0.242 | 0.113 | 0.445 |  |
| Subsp. transition | 0.0511 | 0.0200 | 0.125 |  |
| Subsp. *pauciflora* | 0.0000265 | 0.00000197 | 0.000355 |  |

PCA analysis of trait associations demonstrated clustering of variation within subspecies (Figure **4**) as well as clustering within subspecies irrespective of mean site elevations (Appendix S1: Figure S**3**). The first three PCA dimensions (explaining 58% of total variance) were influenced by anatomically-derived wood traits (*Kv*, LF, *K*s and VD) and tissue properties (FD, WD_branch_, RBT_bole_) as well as leaf size, suggesting that clustering within subsp. arises from both continuous variation associated with elevation and discrete differences in traits between subspecies (Figure **3**).

**FIGURE 4.**
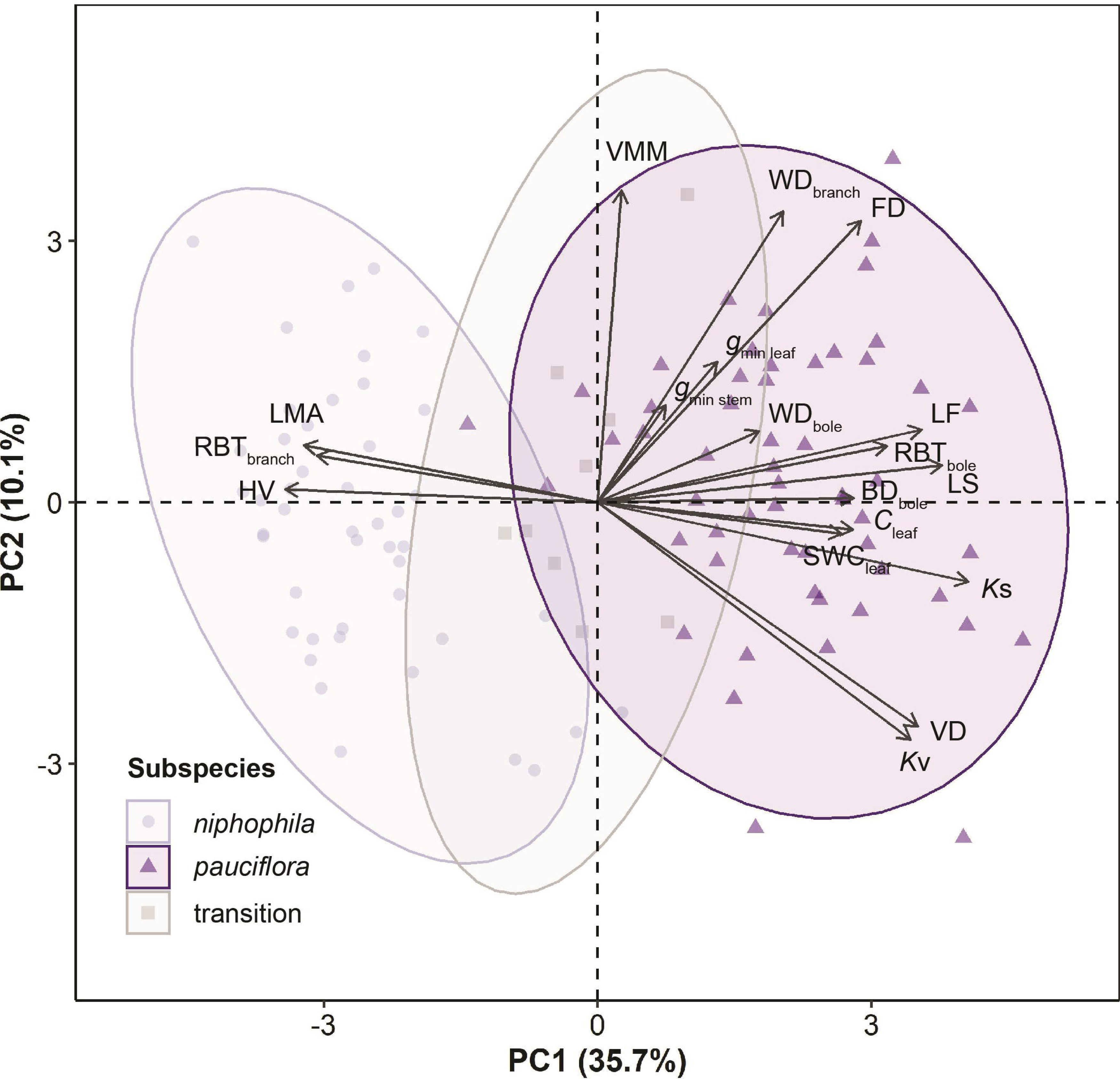
PCA illustrating the clustering of associations between 18 hydraulic and structural traits for two Eucalyptus pauciflora subspecies and the hybridised transition zone between them (n = 111). Symbols: Circles (subsp. niphophila; squares (subsp. transition); triangles (subsp. pauciflora). Abbrev.: *C* (hydraulic capacitance; BD (bark density); FD (fibre density); *g*_min_ (minimum surface conductance to water vapour); HV (Huber value); *K* (hydraulic conductivity of stem (*K*s) and mean vessel (*K*v)); LF (lumen fraction); LMA (leaf mass per unit area); LS (leaf size); RBT (relative bark thickness); SWC (saturated water content); VD (vessel diameter); VMM (vessel density), WD (wood density).

### Associations between trait variation, environmental variation and borer-mediated damage

In a second survey, we explored the direct associations between a subset of traits, bark thickness and wood density and borer-damage severity among dieback-affected and unaffected individuals across a range of elevations, as well as associations with environmental variation. The proportional odds model (Table 1, AIC=1219.1) outperformed a null model (AIC=1547.9, X^2^ =344.88, df =8, P<.00001, *R*^2^_pseudo_=0.42). The probability of severe borer damage differed significantly between subspecies (Figure **5a**, Table **1**, Appendix S1; Table S3). Subspecies *pauciflora* had the lowest mean borer-damage severity score, 0.001 ± 0.001, and very low probabilities of severe borer damage, *Pr.*(severe damage | subsp. *pauciflora*) = 0.00003 (0.00002, 0.00036, CI). Trees in the subspecies transition zone had a higher mean borer-damage severity score of 0.543 ± 0.287 and greater probabilities of severe borer damage than subsp. *pauciflora*, *Pr.*(severe damage | subsp. transition) 0.0512 (0.0200, 0.1250; CI). The highest borer-damage severities were observed within subspecies *niphophila,* with mean borer-damage scores of 1.500 ± 0.334, and the highest probabilities of severe damage, *Pr.*(severe damage | subsp. *niphophila*) = 0.242 (0.113, 0.445, CI).

**FIGURE 5.**
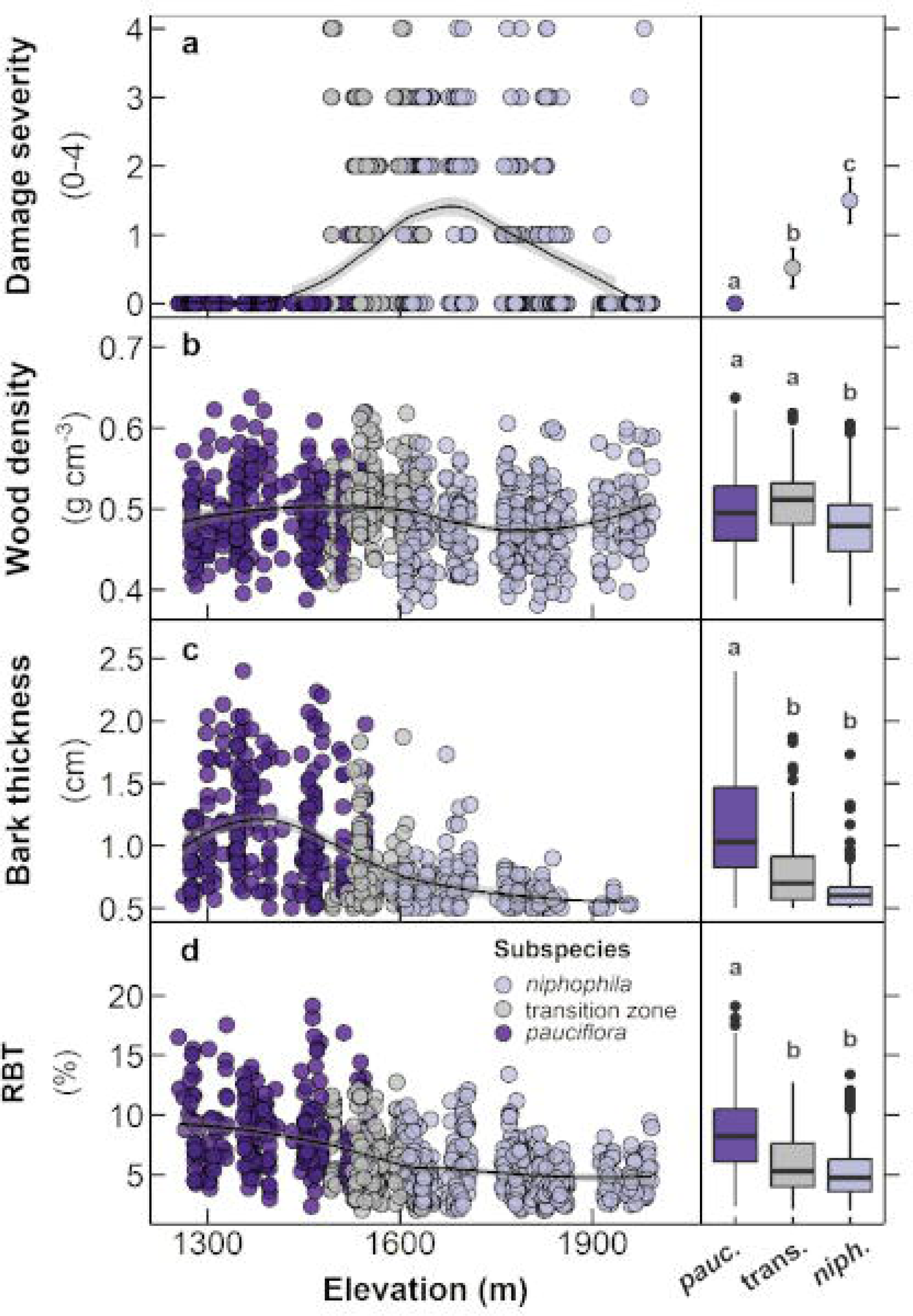
**(a)** Interaction between borer damage severity, subspecies and elevation. An increase in borer damage begins at the subsp. transition zone, peaks at ∼1675 and declines with increasing elevation, resulting in a strong subspecies-elevation interaction (*P*<0.001). **(b)** Variation in wood density across the geographic distributions of subspecies. **(c)** Decline in bark thickness (BT) and **(d)** relative bark thickness (RBT, % of DBH) across elevation, and an increase in borer damage severity. Fitted lines are weighted averages. Significance values were obtained from ordinal regression. RHS insets compare mean subspecies’ responses (*P* <.05).

Between subspecies, we observed mixed interactions between the probability of increased borer-damage severity and elevation (*P* <0.0001, Table 1). Borer-damage severity increased with elevation in both subsp. *pauciflora* (*P* <.0001) and in trees in the transition zone (*P* <.001), and decreased with elevation in subsp. *niphophila* (*P* <.0001, Table 1), resulting in a broad peak in damage severity at ∼1675m (Figure **5a**). Further, 11% of trees surveyed in the transition zone and at higher elevations had dieback severity scores of 1, indicating recent borer activity, i.e. borer-frass holes and wounding stains.

Of the traits included in the borer-damage survey - wood density, absolute bark thickness and relative bark thickness - all differed between subsp. *pauciflora* and subsp. *niphophila;* however, traits of trees of the transition zone were not intermediate but were similar to either subsp. *pauciflora* or subsp. *niphophila,* depending on the trait. Mean wood density was not different between subsp. *pauciflora* and the subsp. transition zone (*P* = 0.22), however, was lower in subsp. *niphophila* relative to both the transition zone (*P*<.001) and subsp. *pauciflora* (*P*=0.03; Figure **5b**). Conversely, absolute bark thickness was higher in subsp. *pauciflora* relative to both the transition zone (*P* <.0001) and subsp. *niphophila* (*P* <.0001), but did not differ between trees of the transition zone and subsp. *niphophila* (*P* = 0.102; Figure **5c**). Similarly, relative bark thickness was higher in subsp. *pauciflora* relative to both the transition zone (*P*<.0001) and subsp. *niphophila* (*P* <.0001), but did not differ between trees of the transition zone and subsp. *niphophila* (*P* = 0.22; Figure **5d**).

Wood density and absolute bark thickness were not associated with variation in the probability of severe borer damage (Figure **6a,b**; *P* = 0.61 and *P* = 0.73, respectively). However, declining RBT (*P* <0.01) and increased DBH (*P* <0.05) were both associated with an increased probability of severe borer damage (Figure **6c**, Table **S3**). Comparison of allometric relationships between RBT and DBH within subspecies revealed reduced bark thicknesses (absolute and relative) across all size classes in subsp. *niphophila* and the transition zone relative to subsp. *pauciflora* (Figure **6d**).

**FIGURE 6.**
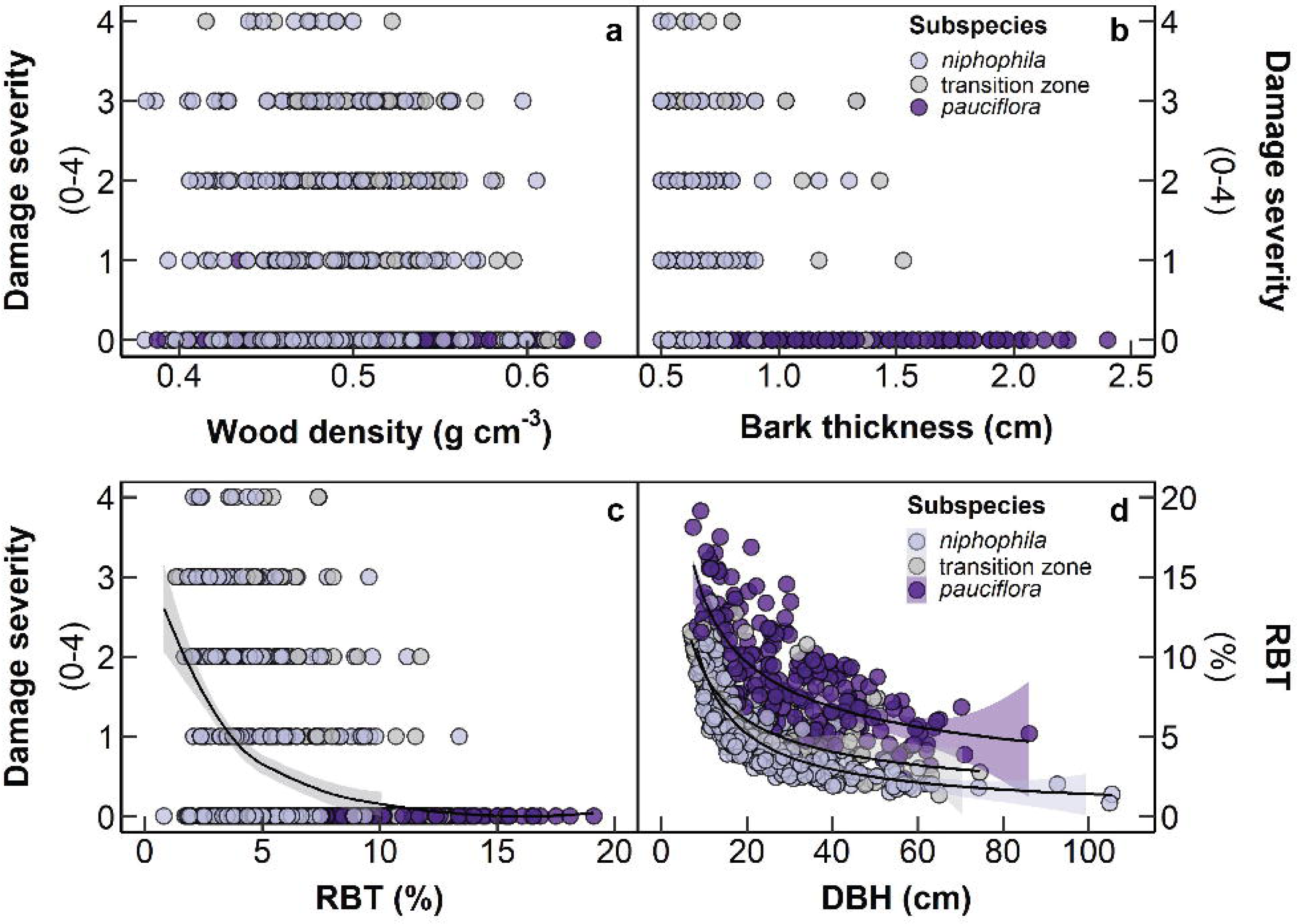
Variation in borer damage severity with (a) wood density (*P*=0.73), (b) absolute bark thickness (RBT, *P*=0.66), and (c) relative bark thickness (*P* < .001). (d) Differences among subspecies in allometric relationships between RBT and diameter at breast height (DBH). Fitted lines are locally weighted averages. Significance values were obtained from ordinal regression, Table 1. *n* = 774 trees.

To explore the decline in borer-damage severity with increasing elevation observed in subsp. *niphophila*, we repeated the above analysis using subsp. *niphophila* data only. Notably, in subsp. *niphophila* greater borer-damage severity variation was not associated with variation in RBT, BD or WD but with increased DBH (*P* = 0.028) and lower elevations (*P*<.0001; Appendix S1: Table **S4a**). Variation in elevation may act as a proxy for many environmental variables ultimately responsible for variation in borer-damage severity. Therefore, we further explored the relative predictive strength of modelled environmental variables highly correlated with elevation for predicting the decline in borer-damage severity with increasing elevation observed in subsp. *niphophila* (*r*^2^>.7, Appendix S1: Table **S5**) via comparison of several models containing single-term elevation substitutions (Appendix S1: Table **S4b**). Increased borer-damage severity was correlated with lower mean summer precipitation (*P*<.01), lower mean annual precipitation (*P*<.05) and lower mean winter precipitation (*P* <.05), and higher mean summer temperatures (*P*<.05), higher mean annual temperatures (*P*<.05), higher mean winter temperatures (*P*<0.05), and higher minimum temperatures (*P*<.05). These bioclimatic variables could be all substituted for elevation with only moderate declines in model performance (Appendix S1: Figure **S4**, Table **S4b**). Notably, elevation was the strongest predictor (*P* <.0001), suggesting that elevation served as a composite variable, integrating multiple contributing bioclimatic predictors.

## DISCUSSION

Increased dieback at intermediate elevations may be driven by spatial variation in environmental conditions that favour borer success at these locations, spatial variation in host-tree suitability favouring borer success at these locations, or a combination of the two. While the relative contributions of these factors are not easy to parse, and both may also have acute and long-term temporal dimensions, the present study provides evidence consistent with the roles of both host-tree suitability and environmental conditions in the moderation of observed elevation-dependent spatial distributions of the current snow-gum dieback occurring in the subalpine woodlands of south-eastern Australia. In the present study, we provide evidence that patterns of dieback severity are associated with differences in specific tissue properties among snow-gum subspecies as well as temperature and precipitation gradients related to elevation. Our initial trait survey identified several traits that varied discreetly between the two snow-gum subspecies and across the subspecies transition zone. The subsequent borer-damage severity survey characterised increased borer damage through the montane-subalpine snow gum subspecies’ transition zone, with a broad peak at borer-damage severity at ∼1675m, corresponding to the lowest elevations of subsp. *niphophila*, followed by a decline in borer-damage severity at the highest elevations. Increased borer-damage severity was associated with lower relative bark thickness, a tissue trait distinctly lower in subsp. *niphophila* relative to subsp. *pauciflora.* The decline in borer-damage risk at the highest elevations was correlated with spatial variation in several environmental variables, notably higher precipitation and lower temperatures. Improved resolution of these complex biotic-environmental interactions will be essential for the reliable modelling of forest vulnerability and effective management for forest resilience under future climates.

### Higher risk of severe borer damage in subspecies niphophila

The probability of borer damage was higher at the lowest elevations of subsp. *niphophila.* This association may suggest that *Phoracantha* preferentially select subsp. *niphophila* for one or more of its traits. While both bark thickness and wood density differed significantly between subspecies, only lower relative bark thickness (RBT) was associated with increased severity of borer damage (Figure **6c**, Table 1). Our findings suggest that differences in bark thickness between subspecies may moderate the differential risk of severe borer damage between subalpine and montane snow-gum subspecies. Greater bark thicknesses in subsp. *pauciflora* were associated with reduced risk of borer damage across their montane distribution; conversely, the decline in bark thickness across the subsp. transition zone was associated with increasingly severe borer damage beginning in the subsp. transition zone and peaking in lower-elevation subsp. *niphophila*. As RBT integrates both absolute bark thickness (BT) and DBH, the highest RBT is observed in smaller stems, while the lowest RBT is observed in larger-diameter stems (Figure **6d**). In other wood-boring species, within-tree distributions of feeding galleries have been shown to follow quadratic relationships in response to variation in bark thickness and stem diameter (Timms, Smith and De Groot, 2006), with gallery height minima and maxima thought to be influenced by spatial variation in bark nutritional quality (Haack, Wilkinson and Foltz, 1987; Hanks, Paine and Millar, 1993; Manville *et al*., 2002), and larval protection from predators, parasites and environmental conditions (Paine *et al*., 2000; Wermelinger, 2002). Preferences of other *Phoracantha* species for larger diameter individuals and lower bole positions (larger diameter) are associated with higher larval survival and emergence, as well as higher phloem densities (Hanks, Paine and Millar, 2005; Nahrung *et al*., 2014; Seaton, Matusick and Hardy, 2020).

### Borer damage risk correlated with spatial variation in temperature and precipitation

One alternate hypothesis for the observed increase in borer damage among low elevation subsp. *niphophila* is that favourable environmental conditions at specific elevations may account for increased borer activity, i.e. a coincidental increase in borer activity at elevations where only subsp. *niphophila* occur. In subsp. *niphophila*, variation in borer-damage severity was negatively correlated with increasing elevation (Figure **5a**, Appendix S1: Table **S4a**), a composite spatial variable integrating variation in numerous thermal and precipitation regimes distinct to subalpine environments (Appendix S1: Table **S4b**, Figure **S2**). Notably, increases in borer-damage severity were correlated with decreases in several precipitation summary statistics (mean summer, winter and annual precipitation) and increases in several temperature summary statistics (mean summer, winter and annual temperature, and minimum temperature; Appendix S1: Table S3b). While these variables covary continuously with elevation, therefore span both subspecies’ distribution, this hypothesis suggests that woodborer activity would be greatest at the lowest elevations due to warmer, drier conditions. As we observed a decline in borer damage severity below the subspecies’ transition zone, our observations suggest that despite favourable environmental conditions at lower elevations, borer success may be limited by a lack of suitable host-trees.

### Drivers of snow-gum dieback

Assuming subsp. *niphophila’s* bark thickness has not changed recently; an open question remains as to the drivers of the recent increase in borer activity. As *Phorocantha* outbreaks often co-occur with drought and heatwave conditions (Hanks, Paine and Millar, 1991; Nahrung *et al*., 2014; Seaton *et al*., 2015; Seaton, Matusick and Hardy, 2020), associations between environmental conditions and borer-dieback severity in the present study also need to be interpreted with respect to variation in environmental conditions over both short and long timescales. The Australian Alpine bioregion is subject to acute El Niño drought events and heat waves, as well as persistent long-term regional warming and drying (Lawrence *et al*., 2022; Verrall *et al*., 2023). Therefore, the persistent extension of the growing season with warming climates might be predicted to increase background *Phorocantha* population pressure at any given elevation, and could extend *Phorocantha’*s thermal niche into higher elevations, as has been reported in other forest systems (Cudmore *et al*., 2010; Pureswaran, Roques and Battisti, 2018). This long-term prediction may also be expected to be further amplified during periodic acute drought events. While future work may be able to characterise temporal variation in environmental conditions associated with snow-gum dieback onset and spread, the present study provides some preliminary insights. Consistent with both of these general predictions, increased wood-borer damage was first observed around 2017 following the extreme El Niño event of 2015/2016 (Santoso, Mcphaden and Cai, 2017). However, the present study, conducted in 2022, also noted 11% of trees surveyed in the transition zone and above demonstrated evidence of recent borer damage. This ongoing presence of borer activity despite the recent easing of El Niño conditions and rehydration of trees may suggest a persistent increase in background borer-pressure, or that the hydraulic dysfunction caused by borer galleries during acute El Niño events may improve the habitat quality for borer larvae rendering successfully colonised trees vulnerable to persistent borer activity following the easing of drought conditions. Another non-mutually exclusive explanation is that recent unprecedented drought stress improved subsp. *niphophila*’s host-tree properties, i.e. by reducing resistance to larvae penetration and/or improving nutritional quality via phloem dehydration. Closer monitoring of seasonal drought stress, tissue water content, and variation in phloem density in borer-sensitive subalpine snow gums will be necessary to resolve the role drought stress as a possible amplifying factor in subalpine *Phoracantha* mediated dieback. Greater resolution of the interaction between environmental variation and the life history characteristics of *Phoracantha mastersii* will be central to understanding ongoing risks to subalpine woodlands, particularly interactions between season length, mean and minimum temperatures and beetle survival and reproductive success. Finally, while the present study focused on only two snow-gum subspecies in one region, the reported presence of borer-mediated dieback in other snow-gum subspecies in the Australian Alps may provide additional opportunities to assess whether trait × environment × dieback relationships are generalisable across subspecies in other subalpine woodlands where conditions, traits and dieback severity may vary.

### Conclusion

Subalpine forests form a critical structural foundation for biodiversity and mountain-derived ecosystem services (Immerzeel *et al*., 2020). The present study indicates that spatial patterns of subalpine forest mortality may depend on interactions between species traits and the dual challenges of changing abiotic and biotic conditions. Cumulatively, our findings are consistent with the conclusion that elevation-dependent patterns of snow-gum dieback may be moderated by both environmental conditions and host-tree traits. We posit that severe dieback occurs were improved environmental conditions for growth of *Phoracantha mastersii* growth intersects with locations where trees possess trait values that are inadequate to resist damage. If accurate, this interpretation suggests that under warmer and drier future climates, subalpine snow-gum forests may be subject to an increased risk of severe borer-mediated forest dieback. For many unmanaged forests, greater resolution of these complex biotic-environmental interactions will be essential for the reliable modelling and effective management for forest resilience under future climates (Crimmins *et al*., 2011; Svenning and Sandel, 2013; Ordonez, Williams and Svenning, 2016).

## Author contributions

CB, HC, ABN, JWJ, and HB planned and designed the research. CB, HC, PC, MD, JD, JDE, DG, RH, JK, AM, AN, YY, OY, and HB performed experiments and conducted fieldwork. CB analysed the data with input from MCB, JB, HC, TIF, ABN, and HB. CB wrote the manuscript with input from all authors.

## Acknowledgements

The authors acknowledge the traditional owners of the lands on which the research was conducted, the Monaro-Ngarigo and Ngunnawal peoples. CB, RJH, HC and JWJ were supported by an Australian Government Research Training Program (RTP) Scholarships. CB was also supported by the Holsworth Wildlife Research Endowment – Equity Trustees Charitable Foundation & the Ecological Society of Australia, as well as the Joyce W. Vickery Scientific Research Fund. HB was supported by an ARC DECRA.

## Data sharing and data Availability statement

The datasets generated and analysed during this study will be made openly available via the ANU Data Commons, upon article acceptance.

## Appendix S1 contents

**Contents:**

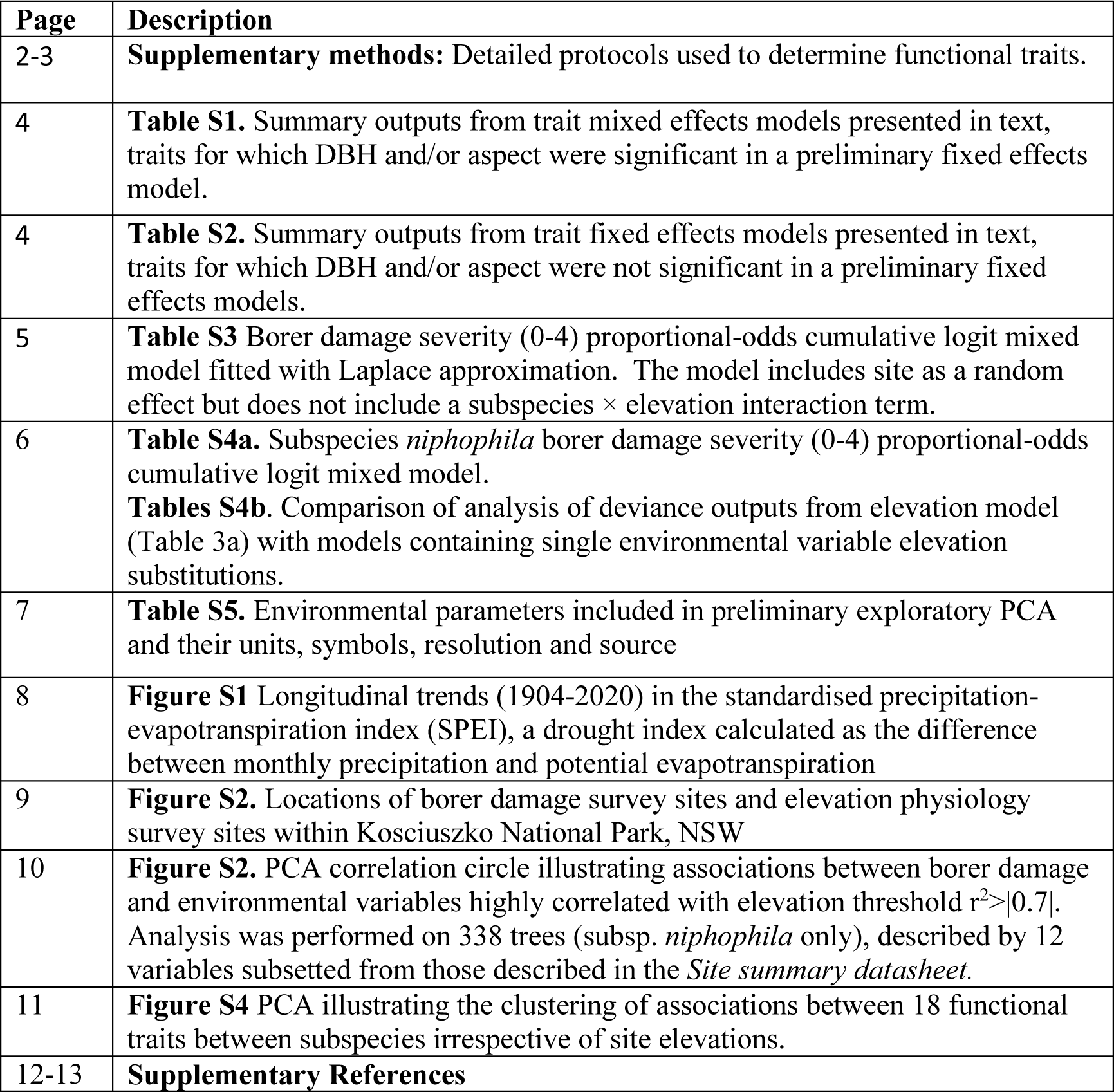

