## Appendix S1 for "Elevation-dependent patterns of snow-gum dieback are associated with subspecies’ trait differences and environmental variation"

**Title:**
Elevation-dependent patterns of snow-gum dieback are associated with subspecies’ trait differences and environmental variation.
**Authors:**
Callum Bryant^1^*, Marilyn C. Ball^1^, Justin Borevitz, Matthew T. Brookhouse^2^, Hannah Carle^1^, Pia Cunningham^1^, Mei Davey^1^, James Davies^1^, Ashleigh Eason^1^, Joseph D. Erskine^1^, Tomas I. Fuenzalida, Dmitry Grishin^1^, Rosalie Harris^1^, Jessica Kriticos^1^, Aaron Midson^2^, Adrienne B. Nicotra^1^, Annabelle Nshuti^1^, Jessica Ward-Jones^2^, Yolanda Yau^1^, Olivia Young^1^, Helen Bothwell^1,3^
**Contact information:**
^1^ Research School of Biology, Australian National University, Canberra, ACT, Australia.
^2^ Fenner School of Environment and Society, Australian National University, Canberra, ACT, Australia.
^3^ Warnell School of Forestry and Natural Resources, University of Georgia, Athens, Georgia, USA

**Contents:**

| **Page** | **Description** |
| --- | --- |
| 2-3 | **Supplementary methods:** Detailed protocols used to determine functional traits. |
| 4 | **Table S1.** Summary outputs from trait mixed effects models presented in text, traits for which DBH and/or aspect were significant in a preliminary fixed effects model. |
| 4 | **Table S2.** Summary outputs from trait fixed effects models presented in text, traits for which DBH and/or aspect were not significant in a preliminary fixed effects models. |
| 5 | **Table S3** Borer damage severity (0-4) proportional-odds cumulative logit mixed model fitted with Laplace approximation. Model includes site as a random effect, but does not include a subspecies: elevation interaction term. |
| 6 | **Table S4a.** Subspecies *niphophila* borer damage severity (0-4) proportional-odds cumulative logit mixed model. **Tables S4b**. Comparison of analysis of deviance outputs from elevation model (Table 3a) with models containing single environmental variable elevation substitutions. |
| 7 | **Table S5.** Environmental parameters included in preliminary exploratory PCA and their units, symbols, resolution and source |
| 8 | **Figure S1** Longitudinal trends (1904-2020) in standardized precipitation-evapotranspiration index (SPEI), a drought index calculated as the difference between monthly precipitation and potential evapotranspiration |
| 9 | **Figure S2.** Locations of borer damage survey sites and elevation physiology survey sites within Kosciuszko National Park, NSW |
| 10 | **Figure S2.** PCA correlation circle illustrating associations between borer damage and environmental variables highly correlated with elevation threshold r^2^>\|0.7\|. Analysis was performed on 338 trees (subsp. *niphophila* only), described by 12 variables subsetted from those described in *Site summary datasheet.* |
| 11 | **Figure S4** PCA illustrating clustering of associations between 18 functional traits between subspecies irrespective of site elevations. |
| 12-13 | **Supplementary References** |

**Supplementary methods:**

*SPEI Index:*

The SPEI index is a regionally normalised drought index, calculated from the difference between monthly totals of precipitation and potential evapotranspiration, with negative values representing increased aridity due to reduced precipitation and/or increased potential evapotranspiration (Vicente-Serrano et al., 2010). To assist visualisation, data presented are a rolling 24-month average (SPEI 48). While a strong decline in SPEI data is evident in the recent decades for which snow cover records were also available (1955-2020, *F*_1,790_ = 527, *P*<.0001; Figure 1h), a longer period of SPEI data is available (1904-2020) capturing periods negative SPEI in the early 20^th^ century, suggesting that the recent decline must be interpreted with caution (Figure S2).

*Protocols used to determine functional traits.*

Tree height was measured using a laser range finder (Nikon Forestry Pro, Nikon Vision Co. Ltd. Tokyo). From each tree, pole pruners were used to sample sun-exposed canopy branches up to 1m long. Branches were transported back to the lab in dark plastic bags humidified with moistened paper. Upon returning to the lab branches were recut under water and allowed to rehydrate under dark conditions, before measurements were conducted. For each branch, mature leaf area and leaf mass per unit area were averaged from ten mature leaves measured using a leaf area meter (Li3100C, Li-COR Biosciences, Nebraska), and dry mass assessed after oven drying as above. Sapwood area: leaf area ratios (Huber values; HV) were assessed in 70cm long branches with methods previously described (Pérez-Harguindeguy *et al.*, 2013; Mencuccini *et al.*, 2019). Briefly, all leaves were excised from branches, and total branch leaf areas were measured as above. Branch diameter was averaged from three triangulated measurements made with digital micrometer (5202, Shahe, Zhejiang, China). Bark was then excised with a razor blade, and sapwood area estimated from the average of three diameter measurements. Branch bark thickness (BT_branch_) was determined by subtracting sapwood diameter from total branch diameter and dividing by two. Bark was then removed from ~3cm segment stem excised at the base of each branch, and sapwood volume and dry mass were measured as above, to determine branch sapwood density (WD_branch_). From the base of each branch, a neighbouring 3cm segment was then placed in 30% ethanol for subsequent analyses of vessel and xylem property traits. DBH was measured as above. Bole bark thickness (BT_bole_) and bole relative bark thickness (RBT_bole_) were determined as above. Bole bark density was determined with aforementioned methods using full thickness bark sample, ~1.5 cm diameter. Bark samples excised at breast height using a bark knife were transport back to the lab wrapped in plastic film. Bole sapwood density (WD_bole_) was measured using 1.5cm x 2mm micro cores sampled at breast height (Trephor, Belluno, Italy), then transported back to the lab in 1.5mL Eppendorf tubes moistened with 10µL of water.

Leaf saturated water content (SWC_leaf_; g_water_ g^-1^_dry mass_ at 0MPa) and hydraulic capacitance (C_leaf_; Δg_water_ g^-1^_dry mass_ MPa^-1^) were determined from one mature leaf from each individual by constructing pressure-volume curves, using the bench drying method (Tyree & Hammel, 1972; Bartlett *et al.*, 2012). Leaf water potentials (Ψ_leaf_) were measured using a Scholander pressure chamber (1050D, PMS Instrument Albany, USA) at intervals of 5 mg decline in fresh mass (XP 205 Metter Toledo balance, Mettler – Toledo – Ltd., Griefense, Switzerland). Similarly, stem saturated water content (SWC_stem_; g_water_ g^-1^_dry mass_ at 0MPa) and hydraulic capacitance (C_stem_; Δg_water_ g^-1^_dry mass_ MPa^-1^)were measured on 60cm terminal shoots. Upon returning to the lab, from each individual five short stem segments (3-4cm in length) of similar diameter (~0.6-1.1cm diameter) were cut from each individual with each segment containing at least three leaves. Segments were then bench dried for a duration sufficient to allow a decline water potential of 0.3 - 0.5 MPa. Before determination of stem fresh mass and water potential, segments were placed in dark zip lock bags for >40 minutes to enable equilibration of water potentials within stem segments and with attached leaves. Stem fresh mass was then measured, and corresponding stem water potential determined from the average of two equilibrated leaves. Stem dry mass was then determined after oven drying as above. For both stems and leaves, hydraulic capacitance (*C*) was determined from the slope of the linear relationship between changing water content per unit dry mass per change in water potential (𝐶 = 𝛥𝑊𝐶 /𝛥𝛹) and saturated water content (SWC; the tissue water content at 0MPa, estimated from the y-intercept of the linear relationship used to determine *C* (Bartlett *et al.*, 2012; Bryant *et al.*, 2021)*.*

Minimum leaf surface water vapour conductance (*g*_min leaf_) was determined for one mature leaf per individual via the mass loss of detached leaves method (Kerstiens, 1996; Duursma *et al.*, 2019). In brief, petioles were sealed with petroleum jelly and then wrapped with para-film, and then leaves were suspended to dry in the lab with pedestal fans used to interrupt leaf boundary layers. Leaves were weighed at 45 min intervals. By plotting water loss versus time, cuticular transpiration was determined for the linear region (r^2^ > 0.995, from a minimum of 6 points per leaf) after initial stomatal closure. Temperature and relative humidity were measured at 10m intervals (BME280; Core Electronics) and logged with a microcontroller (Arduino MEGA 2560; Keyestudio). *g*_min_ was determined as the cuticular transpiration per two sided area per mole fraction difference in water vapour between leaf and air, where air inside the leaf was assumed saturated with water vapour (Pearcy *et al.*, 2000). Minimum stem surface water vapour conductance (g_min stem_) was determined similarly, however on excised straight stem segments of ~10-15cm in length and ~0.7 - 1.1 cm in diameter, with cut ends sealed with petroleum jelly and then wrapped with para-film (Wolfe, 2020). Stem surface area was estimated as 2πrh, with radius averaged from diameter measurements made at both ends of the segment.

Vessel anatomy for each individual was determined in stem segments taken at the base of a 70cm terminal branch. From each segment 4-5 cross sections (~70 µm thick) were made using a sledge microtome (GSL 1, S. Luccinetti, Schenkung Dapples, Zürich, Switzerland). Sections were stained for 10s with toluidine (0.01% wt/vol) then micrographed at 40X using a light microscope (Axiostar plus, Zeiss, Germany) equipped with a digital camera (MD500, The Logical Interface, Illawong, NSW, Australia). Xylem vessel density (vessel *n* mm^-2^) and vessel lumen areas, were surveyed in 497 ± 152 vessels per individual, images were analysed with ImageJ (Schneider *et al.*, 2012). Vessel diameters were estimated using vessel lumen areas to determine diameter of an equivalent circle. Lumen fraction (mm^2^ mm^-2^) and fibre fractions (1-lumen fraction) were calculated by dividing the total xylem lumen area by the total area of section. Fibre density (g cm^-3^) was determined by dividing sapwood density (g cm^-3^) by the fibre fractional area. Hydraulically weighted mean vessel diameters (D_h_) were calculated as previously described (Tyree & Zimmermann, 1971; Steppe & Lemeur, 2007), and used with the Hagan-Poiseuille equation to calculate the hydraulic conductivity of a mean vessel (*K*v; mg m s^-1^ MPa^-1^) as:

*K*v = (π ρ D_h_^4^)/128 η × ΔΨ/l

where ρ is the density of water (kg m^-3^), η is viscosity of water (MPa s^-1^). *K*v was then multiplied by the mean vessel density per transverse sapwood area to determine the stem specific theoretical hydraulic conductivity (*K*s; kg m s^-1^ MPa^-1^) for each individual (Tyree & Ewers, 1991; Steppe & Lemeur, 2007).

| **Table S1.** Summary outputs from trait mixed effects models presented in text, traits for which DBH and/or aspect were significant in a preliminary fixed effects model. Corresponding ANOVA outputs are presented as annotations in Figure 3, models are presented in order of appearance. | | | | | | | | |
| --- | --- | --- | --- | --- | --- | --- | --- | --- |
| **Trait** | ***Fixed effects*** | | | | | ***Random effects*** | | |
|  | **Predictor** | **Estimate** | **SE** | **df** | **Pr** | **Group** | **Variance** | **Std. Dev** |
| Leaf size | (Intercept) | 3.2704 | 4.8920 | 93.5 | 0.505 | DBH | 5.387 | 2.32 |
|  | Elevation | 0.0043 | 0.0027 | 91.3 | 0.57 | Residual | 16.61 | 4.07 |
|  | Subsp. pauciflora | 17.7346 | 1.7341 | 96.5 | <0.0001 |  |  |  |
| LMA | (Intercept) | 209.7466 | 45.7947 | 65.3 | <.00001 | Aspect | 335.8 | 18.32 |
|  | Elevation | 0.0872 | 0.0266 | 70.4 | <.01 | Residual | 1041.6 | 32.27 |
|  | Subsp. pauciflora | -1.4348 | 14.4533 | 95.8 | 0.921 |  |  |  |
| RBT_bole_ | (Intercept) | 4.2455 | 6.7058 | 45.4 | 0.530 | DBH | 17.13 | 4.13 |
|  | Elevation | 0.00494 | 0.0038 | 49.2 | 0.207 | Aspect | 13.14 | 3.62 |
|  | Subsp. pauciflora | 11.5447 | 2.0586 | 50.5 | <0.0001 | Residual | 9.75 | 3.12 |
| Vessel Diameter | Intercept | 40.326589 | 7.437634 | 33.3 | <.00001 | Aspect | 2.72 | 1.65 |
|  | Elevation | 0.001809 | 0.004316 | 28.8 | 0.678 | Residual | 32.93 | 5.74 |
|  | Subsp. pauciflora | 9.185497 | 2.428724 | 56.6 | <.001 |  |  |  |

| **Table S2.** Summary outputs from trait fixed effects models presented in text, traits for which DBH and/or aspect were not significant in a preliminary fixed effects models. Corresponding ANOVA outputs presented as annotations in Figure 3. Models are presented in order of appearance. | | | | |
| --- | --- | --- | --- | --- |
| **Trait** | **Predictor** | **Estimate** | **Std. Error** | **Pr.** |
| Huber value  (Sapwood: leaf area ratio) | (Intercept) | -5.0971 | 2.6125 | 0.053 |
|  | Elevation | 0.0057 | 0.0014 | <.001 |
|  | Subsp. pauciflora | 6.0806 | 2.7667 | 0.030 |
|  | Elevation × subsp. pauciflora | -0.0043 | 0.0016 | 0.009 |
| RBT_branch_ | (Intercept) | 0.7983 | 6.9845 | 0.909 |
|  | Elevation | 0.0105 | 0.0039 | 0.008 |
|  | Subsp. pauciflora | 16.0498 | 7.3816 | 0.031 |
|  | Elevation × subsp. pauciflora | -0.0118 | 0.0043 | 0.008 |
| Vessel specific hydraulic conductivity | (Intercept) | 0.25418296 | 0.258703 | 0.328 |
|  | Elevation | -0.00007040 | 0.000147 | 0.633 |
|  | Subsp. pauciflora | 0.11173451 | 0.273156 | 0.683 |
|  | Elevation × subsp. pauciflora | -0.00000924 | 0.000163 | 0.955 |
| Stem specific hydraulic conductivity | (Intercept) | 1.77455 | 6.83852 | 0.795 |
|  | Elevation | 0.00160 | 0.00388 | 0.680 |
|  | Subsp. pauciflora | 18.20405 | 7.22055 | <.05 |
|  | Elevation × subsp. pauciflora | -0.00918 | 0.00431 | <.05 |

**Table S3.** Borer damage severity (0-4) proportional-odds cumulative logit mixed model fitted with Laplace approximation. Coefficient estimates and SE are log-odds of a shift in dieback severity category (0-4), holding all other predictors constant.

|  | **Borer damage severity proportional odds** | | |
| --- | --- | --- | --- |
| *Predictors* | *Estimates* | *Std. Error* | *P* |
| Elevation | -2.1246 | 0.5112 | **<0.0001** |
| Subsp. *pauciflora* | -9.4007 | 1.5639 | **<0.0001** |
| Subsp. transition | -1.7772 | 0.7475 | **<0.05** |
| DBH | 0.3522 | 0.1738 | **0.042** |
| WD | 0.0555 | 0.1080 | 0.607 |
| BT | -0.0814 | 0.3036 | 0.788 |
| RBT | 0.6534 | 0.2502 | **<0.01** |
| Aspect | 0.07371 | 0.2143 | 0.739 |
| **Threshold coefficients** | | | |
| *Category change* | *Estimate* | *Std. Error* | *Cumulative odds* |
| 0 \| 1 | -0.8164 | 0.4714 | 0.306 |
| 1 \| 2 | -0.0046 | 0.4701 | 0.494 |
| 2 \| 3 | 1.5337 | 0.4786 | 0.822 |
| 3 \| 4 | 3.8512 | 0.5506 | 0.979 |
| **Random effects** |  |  |  |
| *Group* | *Variance* | *Std. Dev* |  |
| Site Id. | 1.654 | 1.286 |  |
| n _Site Id._ | 53 |  |  |
| Observations | 774 |  |  |
| **Estimated subspecies’ borer damage log-odds** | | | |
| *Subspecies* | *Log-odds* | *Std. Error* | *Odds* |
| Subsp. *niphophila* | -1.14 | 1.564 | 0.242 |
| Subsp. transition | -2.92 | 0.748 | 0.051 |
| Subsp. *pauciflora* | -10.54 | 1.320 | <0.0001 |

**Table S4a.** Subspecies *niphophila* borer damage severity (0-4) proportional-odds cumulative logit mixed model fitted with Laplace approximation. Coefficient estimates and SE are log-odds of a shift in dieback severity category (0-4). *P* values were obtained from likelihood ratio tests of explanatory variables while controlling for remaining predictors.

|  | **Borer damage severity proportional odds** | | | | | |
| --- | --- | --- | --- | --- | --- | --- |
| *Predictors* | *Estimates* | | *Std. Error* | | *P (ANODE)* | |
| Elevation | -1.5663 | | 0.3611 | | **<.0001** | |
| DBH | 0.4651 | | 0.2116 | | **0.028** | |
| WD | 0.1755 | | 0.1256 | | 0.162 | |
| BT | -0.1769 | | 0.1925 | | 0.353 | |
| RBT | -0.3002 | | 0.2385 | | 0.204 | |
| Aspect | 0.0220 | | 0.2810 | | 0.937 | |
| **Threshold coefficients** | | | | | | |
| *Category change* | *Estimate* | | *Std. Error* | |  | |
| 0 \| 1 | 0.572 | | 0.342 | |  | |
| 1 \| 2 | 1.638 | | 0.354 | |  | |
| 2 \| 3 | 3.699 | | 0.404 | |  | |
| 3 \| 4 | 6.159 | | 0.565 | |  | |
| **Random effects** |  | |  | |  | |
| *Group* | *Variance* | | *Std. Dev* | |  | |
| Site Id. | 2.264 | | 1.505 | |  | |
| n _Site Id._ | 27 | |  | |  | |
| Observations | 388 | |  | |  | |
| **Tables S4b**. Comparison of analysis of deviance outputs from elevation model (Table 3a) with models containing single environmental variable elevation substitutions. | | | | | | |
| *Elevation substitution* | | *Variable P* | | *Model AIC* | | *Likelihood ratio statistic* |
| (-) Elevation | | **<.0001** | | 725.84 | | - |
| (-) Mean summer precip. | | **0.0017** | | 731.44 | | -5.601 |
| (-) Mean annual precip. | | **0.0086** | | 734.36 | | -8.522 |
| (+) Mean summer temp. | | **0.0136** | | 735.18 | | -9.341 |
| (+) Mean annual temp | | **0.0199** | | 735.85 | | -10.008 |
| (+) Mean winter temp. | | **0.0221** | | 736.03 | | -10.188 |
| (+) Minimum temp. | | **0.0279** | | 736.43 | | -10.589 |
| (-) Mean winter precip. | | **0.0351** | | 736.83 | | -10.984 |
| Soil Organic Content | | 0.1069 | | 738.67 | | -12.283 |
| Annual Temp Range | | 0.9399 | | 741.28 | | -15.416 |

| **Table S5.** Modelled bioclimatic variables highly correlated with elevation (*r*^2^ = <\|0.7\|), their units, spatial resolution and source. | | | | |
| --- | --- | --- | --- | --- |
| **Environmental variable** | **Units** | **Abbrev.** | **Resolution** | **Source** |
| BIO1 = Annual Mean Temperature | °C | MAT | 1km | WorldClim 2.1 (Fick & Hijmans, 2017) |
| BIO5 = Max Temperature of Warmest Month | °C | Max. temp. |  |  |
| BIO6 = Min Temperature of Coldest Month | °C | Min. temp. |  |  |
| BIO7 = Annual Temperature Range (BIO5-BIO6) | °C | - |  |  |
| BIO8/BIO11 = Mean Temperature of Wettest/ Coldest Quarter | °C | Winter temp. |  |  |
| BIO9/BIO10 = Mean Temperature of Driest/Warmest Quarter | °C | Summer temp. |  |  |
| BIO12 = Annual Precipitation | Mm | MAP |  |  |
| BIO16/BIO19 = Precipitation of Wettest/ Coldest Quarter (Winter) | Mm | Winter prec. |  |  |
| BIO17/BIO18 = Precipitation of driest/ warmest quarter (Summer) | Mm | Summer pre. |  |  |
| Soil Organic Content | Tons/hectare | - | 250m | Soil Grids 250m global dataset (Hengl *et al.*, 2017) |

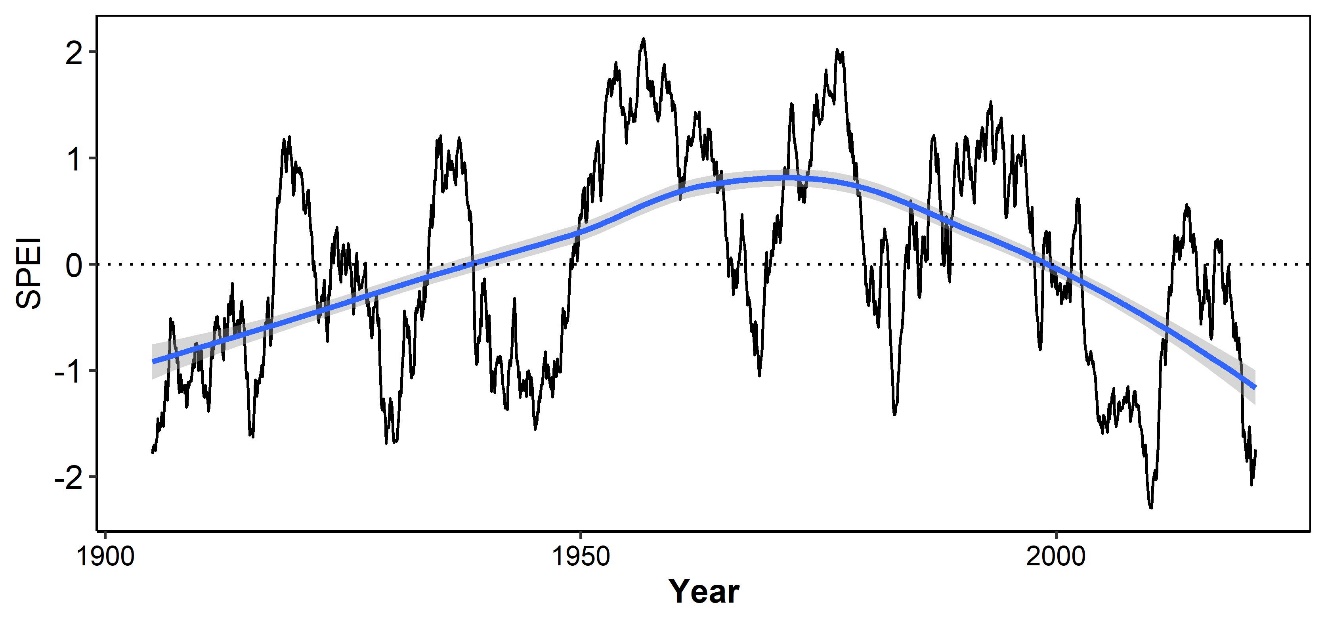

**Figure S1** Longitudinal trends (1904-2020) in standardized precipitation-evapotranspiration index (SPEI), a drought index calculated as the difference between monthly precipitation and potential evapotranspiration (Vicente-Serrano et al. 2010) While monthly SPEI data are available, data presented are SPEI 24, a 24 month rolling SPEI average. Fitted trend line depicts locally weighted averages.

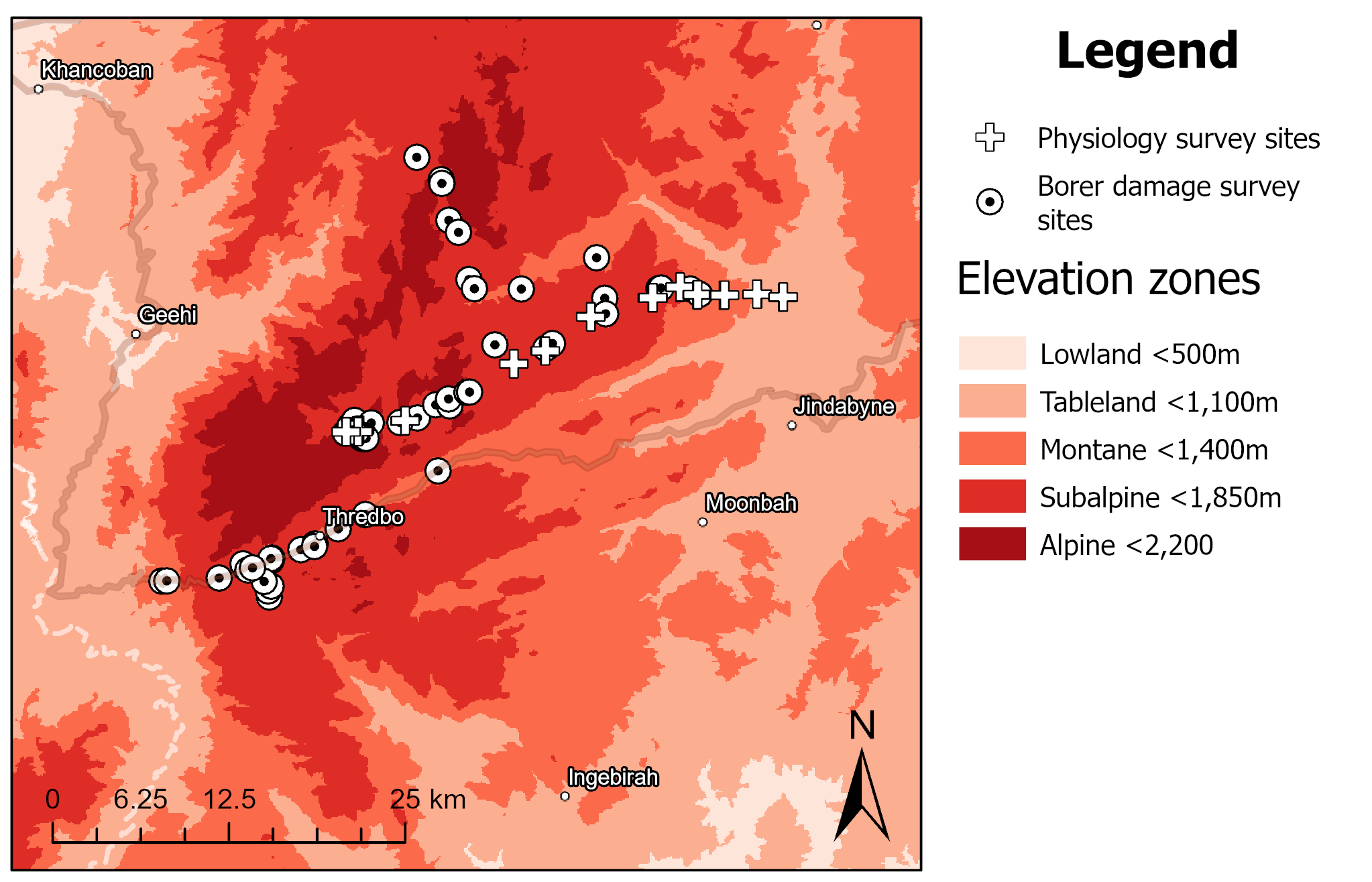

**Figure S2** Locations of borer damage survey sites (circles, *n* = 53) and elevation physiology survey sites (crosses, *n* = 12) within Kosciuszko National Park, NSW. Site details are summarized in Supporting Tables 1 and 2.

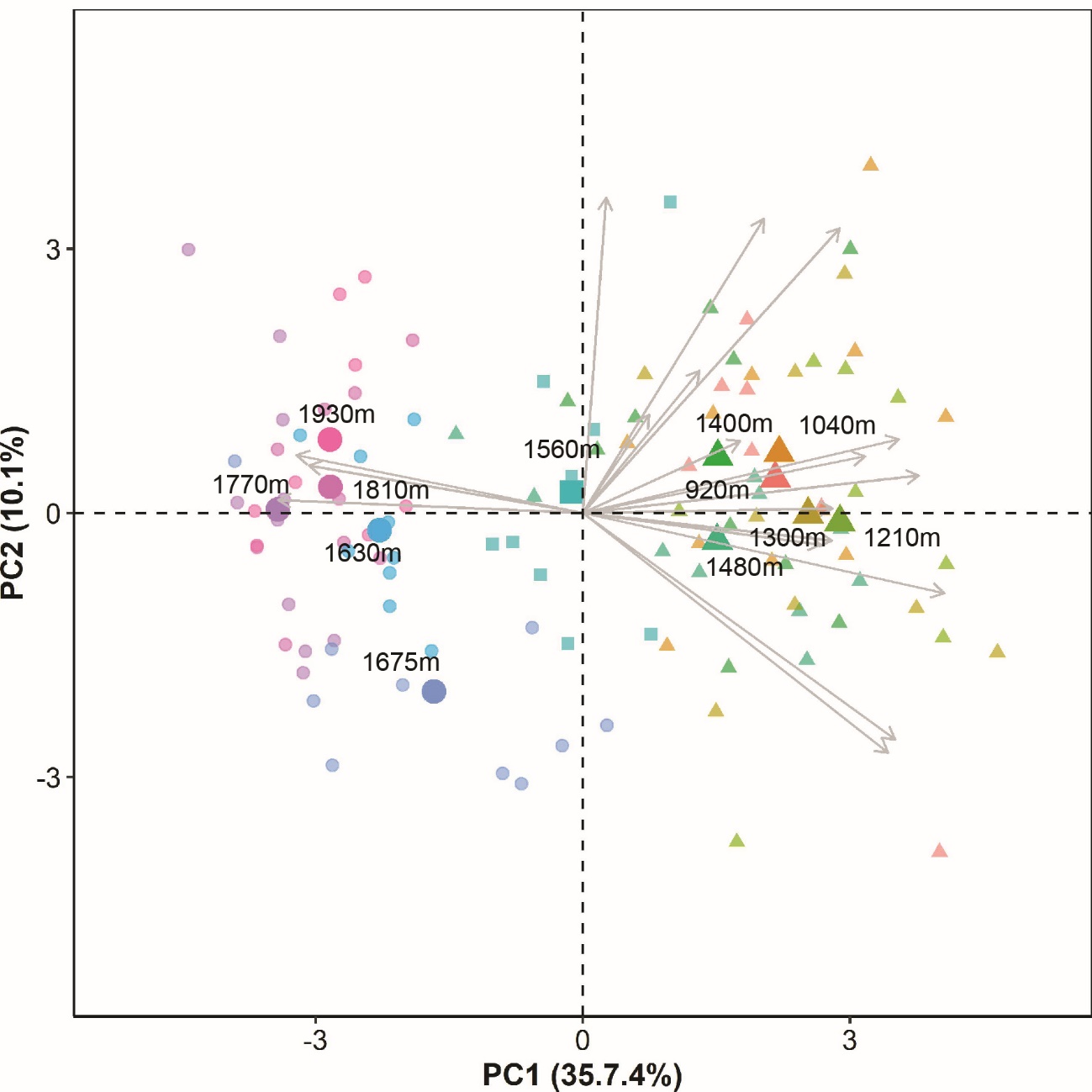
**Figure S3** PCA illustrating clustering of associations between 18 functional traits between subspecies irrespective of site elevations, as shown coloured by subspecies in Figure 4. Symbols: circles (subspecies *niphophila*); square (transition zone); and triangles (subsp. *pauciflora*). Analysis was performed on 111 individuals.

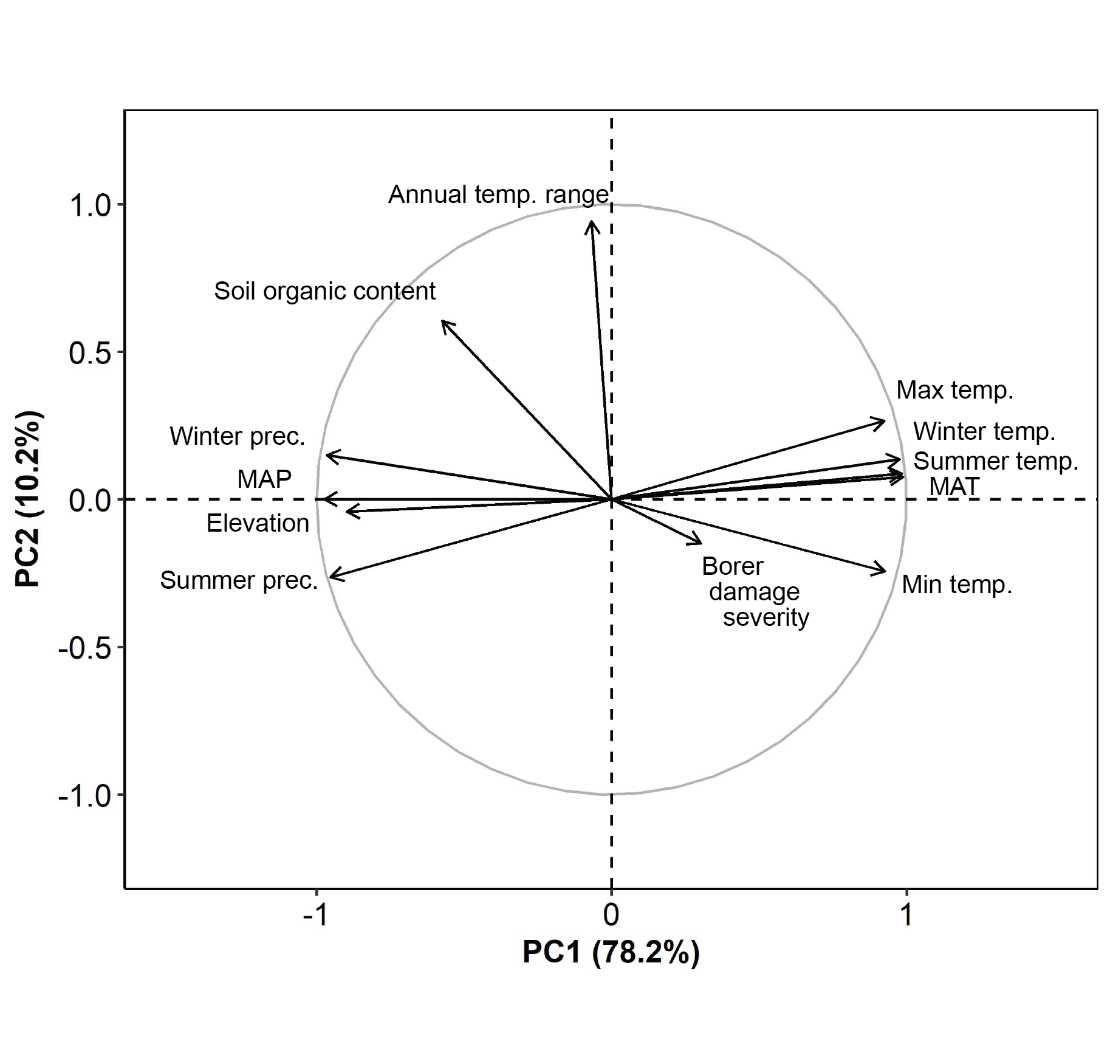

**Figure S4** PCA correlation circle illustrating associations between borer damage and environmental variables highly correlated with elevation, threshold r^2^>|0.7|. Analysis was performed on 338 trees (subsp. *niphophila* only), described by 12 variables subsetted from those described in *SI Table 1*. In interactions between borer damage predictions and elevation for subsp. *niphophila* (Table 2), elevation performs as a proxy composite variable, comprising variation in numerous co-varying environmental parameters.
